# On the intersection between data quality and dynamical modelling of large-scale fMRI signals

**DOI:** 10.1101/2021.05.23.445373

**Authors:** Kevin M. Aquino, Ben Fulcher, Stuart Oldham, Linden Parkes, Leonardo Gollo, Gustavo Deco, Alex Fornito

## Abstract

Large-scale dynamics of the brain are routinely modelled using systems of nonlinear dynamical equations that describe the evolution of population-level activity, with distinct neural populations often coupled according to an empirically measured structural connection matrix. This modelling approach has been used to generate insights into the neural underpinnings of spontaneous brain dynamics, as recorded with techniques such as resting state functional MRI (fMRI). In fMRI, researchers have many degrees of freedom in the way that they can process the data and recent evidence indicates that the choice of pre-processing steps can have a major effect on empirical estimates of functional connectivity. However, the potential influence of such variations on modelling results are seldom considered. Here we show, using three popular whole-brain dynamical models, that different choices during fMRI preprocessing can dramatically affect model fits and interpretations of findings. Critically, we show that the ability of these models to accurately capture patterns in fMRI dynamics is mostly driven by the degree to which they fit global signals rather than interesting sources of coordinated neural dynamics. We show that widespread deflections can arise from simple global synchronisation. We introduce a simple two-parameter model that captures these fluctuations and which performs just as well as more complex, multi-parameter biophysical models. From our combined analyses of data and simulations, we describe benchmarks to evaluate model fit and validity. Although most models are not resilient to denoising, we show that relaxing the approximation of homogeneous neural populations by more explicitly modelling inter-regional effective connectivity can improve model accuracy at the expense of increased model complexity. Our results suggest that many complex biophysical models may be fitting relatively trivial properties of the data, and underscore a need for tighter integration between data quality assurance and model development.

## Introduction

Since the seminal work of Biswal et al. (1), a great deal of research in functional magnetic resonance imaging (fMRI) has focused on understanding spontaneous blood-oxygenation-level-dependent (BOLD) signal fluctuations recorded in the absence of an explicit task––the so-called resting state (2, 3). These spontaneous signal fluctuations are synchronized across distributed neural systems (4) in a way that can robustly reveal, across individuals and species, an underlying network architecture that mirrors task-based co-activation patterns and underlying anatomical connectivity (5–11). These resting-state networks are sufficiently unique to enable identification of individual people, akin to a “connectome fingerprint” (12, 13), and they can be used to predict performance in independently measured behavioural tasks (14, 15). They are heritable (16, 17), influence task-evoked activation and behaviour (18, 19), and are altered in diverse clinical disorders (2, 20), suggesting that they may represent viable clinical biomarkers (21, 22).

These intriguing empirical properties of spontaneous fMRI dynamics raise questions about their physiological origins. Due to limitations on spatial and temporal resolution, fMRI alone cannot reveal these origins. To this end, large-scale biophysical models of neural activity have been used to bridge the gap between the limited measures provided by fMRI and the underling mechanisms that might generate the observed signal properties. The most popular class of dynamic neural models (DNMs) consider activity at the level of mesoscopic neuronal populations, delineated using a particular regional parcellation of the brain (23). The models simulate the aggregate activity of each parcellated region through a series of dynamical equations whose evolution is governed by key biophysical constraints and a pre-specified form of coupling between populations. Regional signal properties, or measures of pairwise or multivariate coupling, can then be compared between the simulated and empirical data. These DMNs allow investigators to artificially manipulate neural populations, test how changes in structural connectivity affect function (7, 8), and make predictions about the effects of lesions on resting-state networks (24).

For present purposes, DNMs have three critical ingredients. The first ingredient is a biophysically relevant model of population dynamics, which describes the net neuronal activity in each brain region. Numerous models and methods have been described (see Deco et al. (23) and Breakspear (25) for reviews) but the key property of all such models is that they assign, to small patches of neural tissue, populations of excitatory and inhibitory neurons that govern the local dynamics. Regardless of the model, coupling between these patches of cortex - represented as nodes in the network – is mediated via excitatory to excitatory projections (though they can be excitatory to inhibitory see Deco et al. (26)). At each node, the resultant neural dynamics are then translated to a BOLD forward model that allows a direct simulation of resting state fMRI (27–29).

The second ingredient of DNMs is an anatomically defined connectivity matrix that couples the neural populations. This matrix is typically defined using empirical estimates of structural connectivity (SC) between parcellated regions (a connectome - see Fig. 1). In human neuroimaging experiments, the connectome is normally mapped using diffusion weighted MRI, so the underlying SC matrix is weighted and undirected (although some models of species with available tract-tracing data have incorporated directionality; e.g,. (30, 31) and see Kale et al. (32) for the importance in using directed connectomes).

**Fig. 1.**
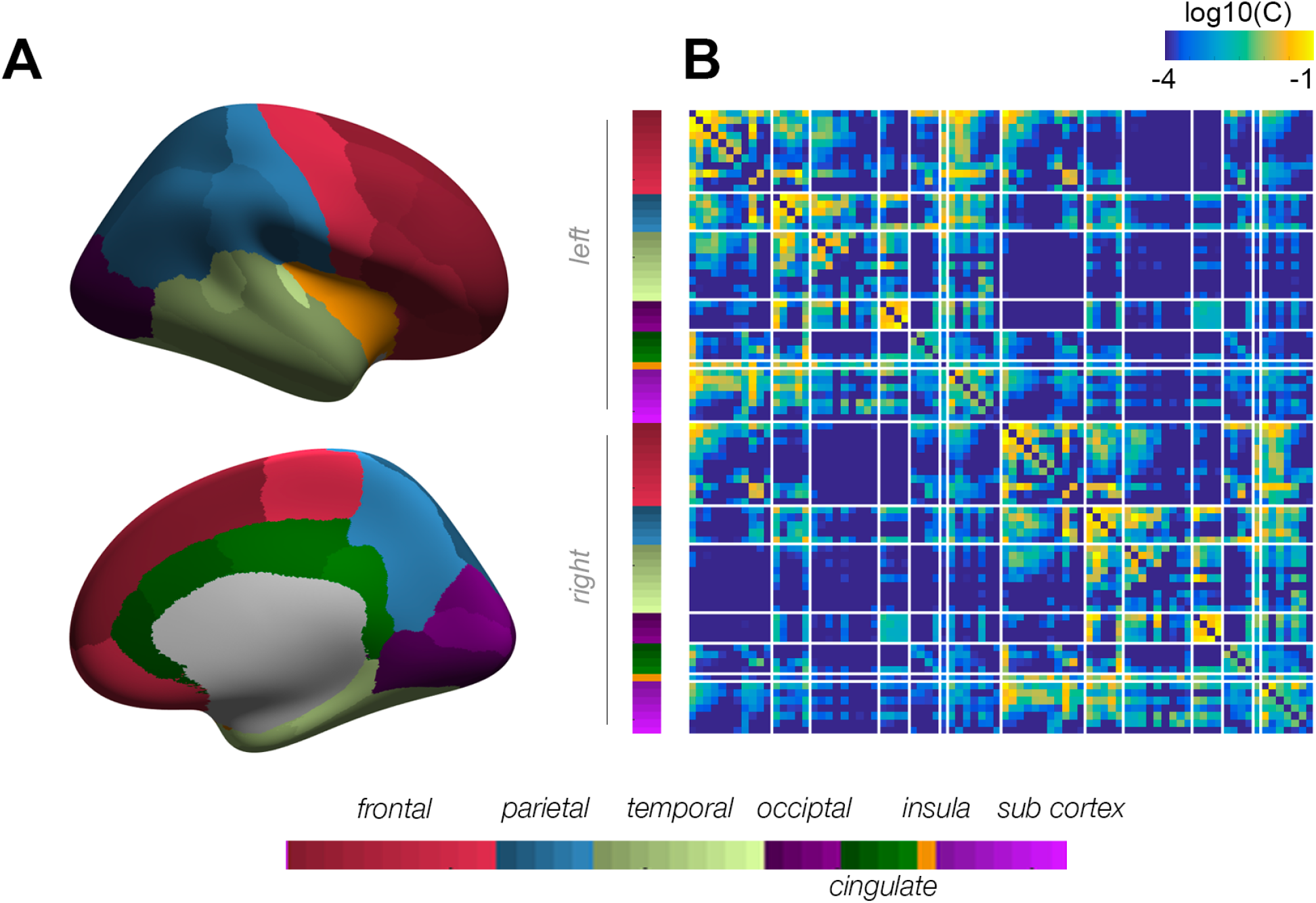
Anatomical parcellation and estimated structural connectivity matrix. A The Desikan killany(59) parcellation displayed on the right inflated hemisphere of the average subject (fsaverage). Colours on the cortex represent lobar regions described below with the shadings of these major regions indicating individual regions. B The estimated group-representative structural connectivity matrix estimated using diffusion weighted imaging (see Methods) with connection weights plotted on a logarithmic scale. The white lines indicate the border of the broad regions across the two cortical hemispheres and the two sides of each region in subcortex.

The third key ingredient of a DNM is its set of biophysically relevant parameters, which can be classed into parameters that are defined globally or at the level of each network node. Node-level parameters translate neuronal dynamics to the population level. For example, the membrane threshold potential for a neuronal population is parametrised by the mean and variance of this voltage potential across a small patch of cortex (23, 26, 33, 34). In most cases, these are fixed at specific values, and all brain regions are treated as having a homogeneous structure and dynamics (8, 26, 35). Recent evidence has suggested benefits to incorporating regional heterogeneity in some of these parameters (31, 34, 36–38) at the cost of more free parameters and increased model complexity.

Most DNMs also contain a global parameter, denoted *G*, that is used to scale the SC matrix uniformly.This is because the weights of the SC matrix are defined in arbitrary units that are not calibrated with the population-level models. The parameter *G* thus defines a baseline level of coupling between populations, and its specific value can have a large influence on the dynamics - values that are too high result in globally synchronous activity and values that are too low cause asynchronous fluctuations (e.g., *G* = 0 are truly uncoupled nodes). It is thus customary to fit the optimal value of *G* to the data, depending on the specific feature one wants to model. Most commonly, *G* is tuned to maximise the similarity between the observed and synthetic FC matrices, but other properties that capture aspects of FC dynamics through time have also been used (34, 36) (we evaluate an example below). We note that there multiple ways of formulating whole-brain models at the population level, from neural masses to fields (23, 39), but here we focus on the most common network-based approach of modelling whole-brain dynamics in which population masses are coupled via the connectome, with a global coupling parameter, *G*.

In any modelling exercise, the quality of the conclusions that one draws from the model depend heavily on the quality of the data being modelled; the old adage “garbage in, garbage out” rings true. fMRI data are notoriously noisy, and resting-state fMRI in particular has been the subject of considerable debate over how various sources of noise should be modelled and removed (40, 41). One issue that has attracted considerable attention is the presence of so-called global signal fluctuations or anatomically widespread signal deflections (WSDs): transient yet coordinated signal changes that affect the vast majority of voxels in an fMRI dataset (42, 43). Such deflections have been tied to various sources of noise, including head motion (42), heart rate variability (42), and respiration (44). Despite several efforts to remove these effects through physiological modelling (45), improved motion correction algorithms, and advances in fMRI acquisition (e.g. (46)) and de-noising (42, 44, 47), they have remained a challenge for the field. Different pipelines show variable efficacy in removing WSDs and other noise sources, and there is no widely-accepted gold standard method (40–42, 44).

WSDs are most clearly visualised using “carpet plots” (or gray plots), in which brain voxels are depicted as rows, time points as columns, and signal changes as grayscale colour variations (43, 48, 49). For example, Figure 2 shows clear WSDs due to head motion (red shaded), whereas Figure 3 shows WSDs unrelated to head motion (blue shaded) which may be tied to respiratory variations (42). These WSDs will obviously increase the global coherence of voxel-level signals, often in a way that increases correlations between FC estimates and measures of noise related to head motion (40– 43) and physiology (42, 48). Such events have important implications for DNMs, given that *G*, which defines the global coupling of the model, is explicitly fitted to the data. More specifically, given a sufficiently noisy dataset, it is likely that the process of optimising *G* in a given DNM will simply fit the WSDs in the data. Such issues of data quality have seldom been considered in the modelling literature, with many DNMs fitted to data processed using a pipeline that does not explicitly include a step (such as GSR for instance) to remove WSDs (26, 34, 50). Thus, it is possible that considerable structured, global noise may still be present in the data.

**Fig. 2.**
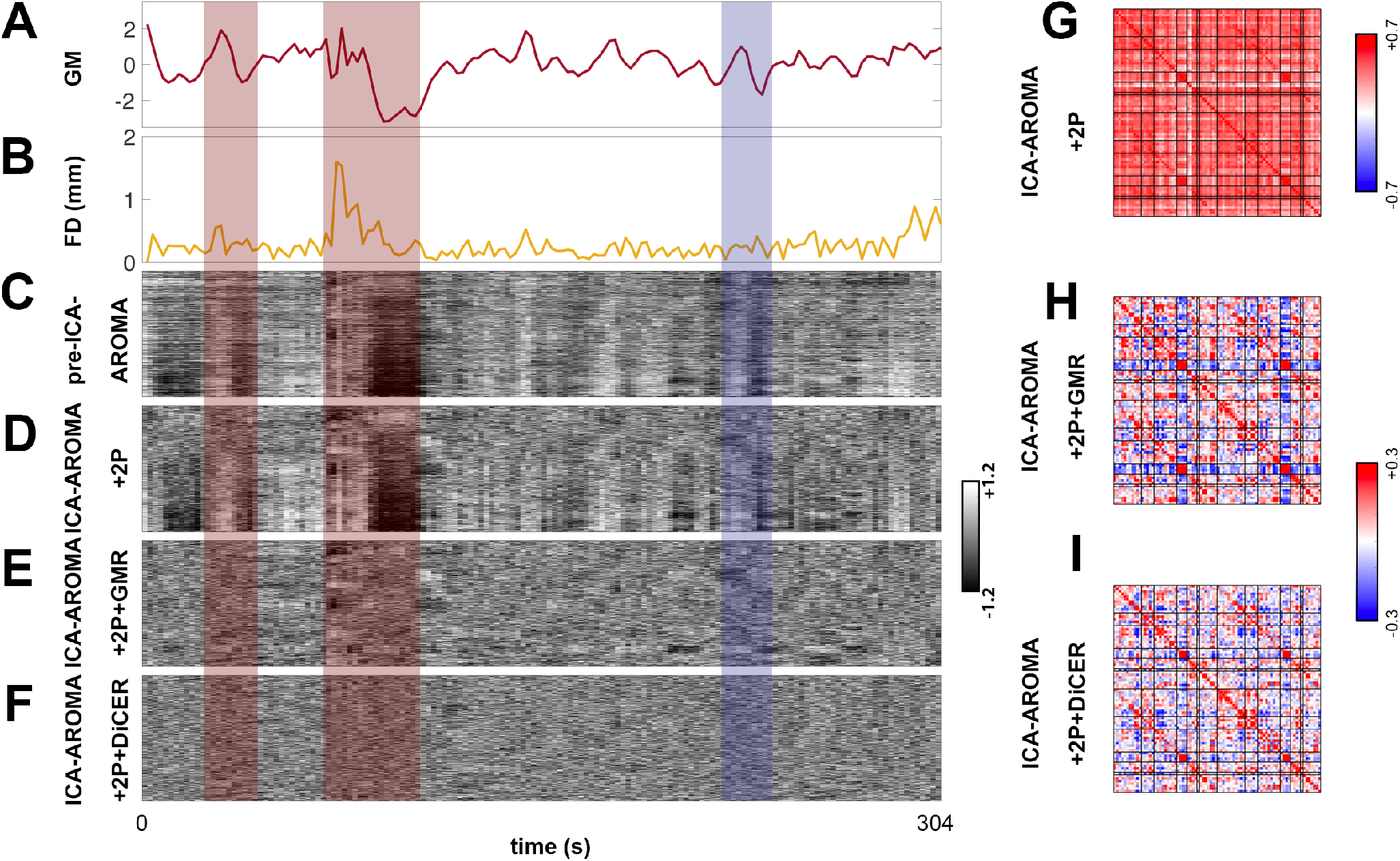
Motion-induced WSDs in subject sub-10290 of the UCLA cohort. (A) The average grey matter signal used for GMR. (B) Framewise displacement (FD) estimate through time. The next four panels are carpet plots for the (C) pre-ICA-AROMA base preprocessing; (D) ICA-AROMA with white matter and CSF regression (+2P) pipeline; (E) Grey matter regression (GMR) following ICA-AROMA+2P pipeline; and (F) Diffuse cluster estimation and regression (DiCER) following ICA-AROMA+2P pipeline. In panels (A)–(F) the red transparent rectangles indicate periods of motion induced WSDs, whereas the blue rectangle indicates WSDs generated without strong motion events. These carpet plots are ordered with respect to cluster ordering (CO) visualisation. Panels G, H, and I show the corresponding FC matrices for ICA-AROMA+2P, ICA-AROMA+2P+GMR, and ICA-AROMA+2P+DiCER, respectively

**Fig. 3.**
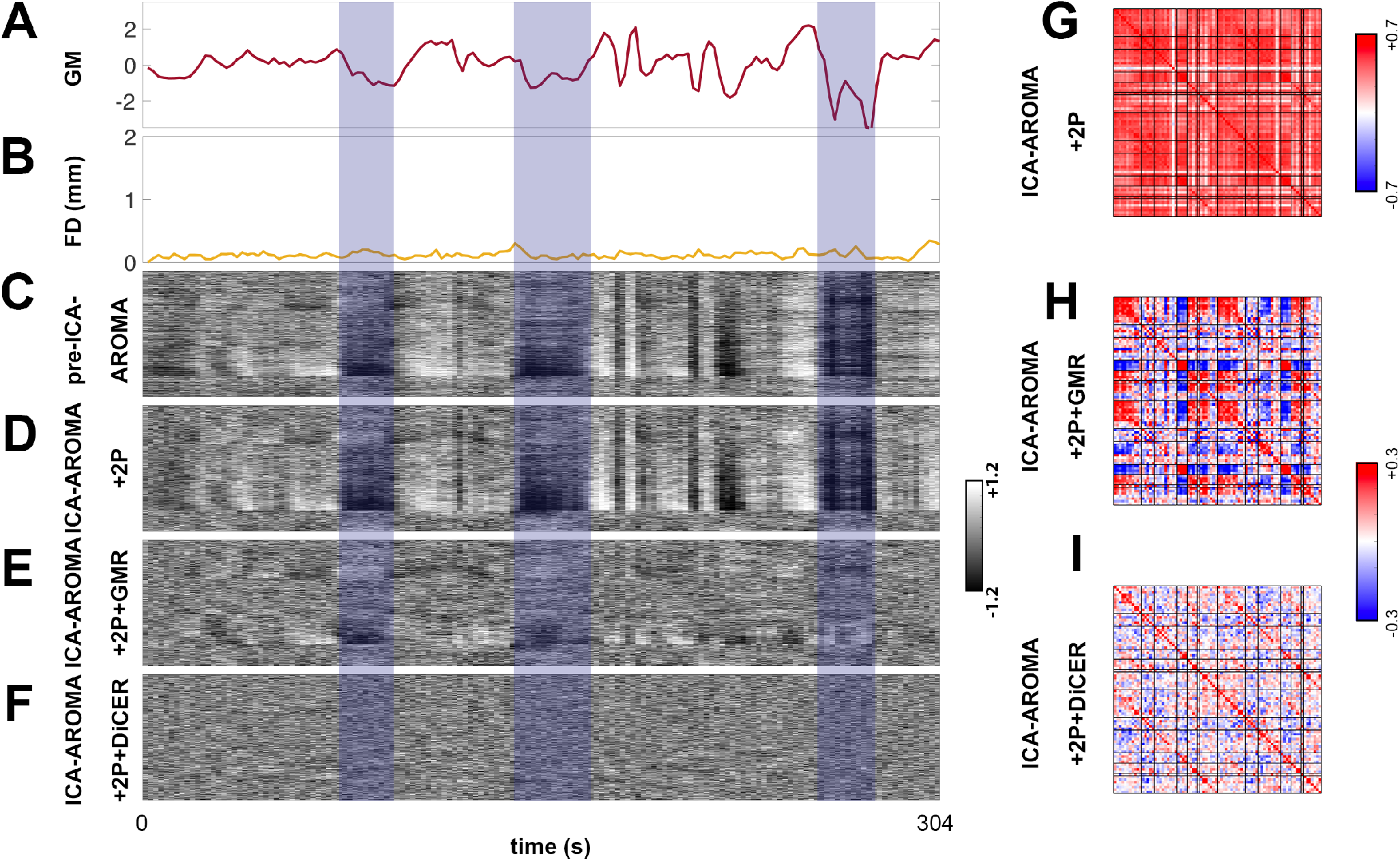
Non-motion-induced WSDs in subject sub-10274 in the UCLA cohort. (A) The average grey matter signal used for GMR. (B) Framewise displacement (FD) estimate through time. The next four panels are carpet plots for the (C) pre-ICA-AROMA base preprocessing; (D) ICA-AROMA with white matter and CSF regression (+2P) pipeline; (E) Grey matter regression (GMR) following ICA-AROMA+2P pipeline; and (F) Diffuse cluster estimation and regression (DiCER) following ICA-AROMA+2P pipeline. In panels (A)–(F) the blue rectangle indicates WSDs generated without strong motion events. These carpet plots are ordered with respect to cluster ordering (CO) visualisation. Panels G, H, and I show the corresponding FC matrices for ICA-AROMA+2P, ICA-AROMA+2P+GMR, and ICA-AROMA+2P+DiCER, respectively.

Previous studies have used DMNs to model large scale brain dynamics and have reproduced FC matrices from structural connectivity patterns across many different experimental cohorts (8, 25, 26, 35, 36, 50–52). Reproducing FC is the baseline measure of model accuracy, and studies have reached relatively high accuracy reaching up to a correlation of *r* = 0.78 (34) between the empirical and the modelled FC matrix. However, reproducing FC structure of data containing globally synchronized activity is relatively simple to reproduce (See results) and many fMRI datasets are well known to contain large WSDs (42, 43) which may be driven by spurious sources (42, 44). This puts into question whether DNMs are really performing well at capturing subtle whole-brain communication dynamics or whether they are simply performing well at capturing relatively trivial globally synchronized WSDs.

In light of these observations, this work investigates the ways in which different denoising strategies (to remove WSDs) in resting-state fMRI data influence DNM fits, and the dynamic regimes that the fitted DNMs display. Specifically, we use a standard open source data set (53) that has previously been denoised using three different approaches (43), which range from “lenient” to “moderate” to “aggressive” in their approach to WSD removal. We test the validity and the fitting quality of three popular models in the literature given the same input data, but denoised in different ways.

We report several key findings. First, like fMRI data, the model simulations themselves exhibit WSDs with varying degrees of strength and brain-wide coverage (i.e., from groups of correlated regions, to whole brain synchronisation). In all these models, the prominence of WSDs tracks the value of the global coupling parameter, *G*. Second, model fits tend to be higher for data with more prominent WSDs, pushing the models into a more globally synchronous regime. Third, with more aggressive preprocessing, model fits reduce and the dynamic regimes of the models transition from globally synchronous to weakly synchronous. Fourth, data that have not been processed to remove large WSDs can be accurately fitted with a two-parameter global signal model that relies only on node degree in the SC matrix, coupled with global and local white noise. This result suggests that large WSDs can be generated through simple mechanisms and that models fitted to data with WSD will only be capturing relatively trivial global synchronization events. Finally, we find that heterogeneous variants of DNMs, in which node and edge parameters are allowed to vary, can improve model fits in more aggressively denoised data, at the expense of increased model complexity. These results highlight the need for careful evaluation of data quality during model-fitting, and a careful consideration of how model fits are interpreted, and a tighter integration between research efforts directed towards data collection and modelling. All analysis code used here is provided as an open source toolbox available at http://github.com/KevinAquino/modelling_comparisons/.

## Imaging Methods

### Resting state fMRI

Here, we used a transparent, open rsfMRI dataset processed using open-source pipelines from `fmriprep` (54).

#### Imaging data & preprocessing

We utilized the open source dataset from the healthy controls of the UCLA Consortium for Neuropsychiatric Phenomics LA5c Study (53) (v00016 openneuro.org/datasets/ds000030/).

The scanning parameters for these data are described in (54). Briefly, task-free fMRI was acquired with an echo-planar imaging (EPI) sequence at 3 T on a Siemens Trio 3T MRI Scanner and a 32-channel receive-only head coil. The fMRI imaging parameters involved echo time (TE) = 30 ms, 3 mm Inplane resolution, 34 slices with 4 mm slice thickness, FA= 90, field of view = 192 mm, matrix = 64 *×* 64, using an oblique and interleaved Gradient echo EPI sequences, TR = 2 s, and a total of 152 Volumes resulting in 304 s of resting state fMRI. A total of 270 subjects were acquired, for which we only focused on 121 healthy controls within the sample.

A “Base” preprocessing pipeline was used on the UCLA dataset using `fmriprep` v1.1.1 as described in (43). Briefly, this involves processing the T1-weighted anatomical images using bias field correction, brain extraction, freesurfer segmentation, and volume normalisation to the MNI 152 Nonlinear Asymmetric template version 2009c (55). For the functional MRI data, the minimal preprocessing steps involve: slice time correction, motion correction, distortion correction using a template based *B*_0_ image, and co-registration to the T1 anatomical image. The code to run all of these analyses is located at https://github.com/BMHLab/DiCER.git and specific details of the algorithms are detailed in Aquino et al. (43). In the following sections, we describe de-noising strategies that follow base preprocessing.

#### Denoising strategies

Functional MRI data were analysed within the MNI 152 Asymmetric 2009c space, resampled to the native BOLD image dimensions of each individual. We resampled any requisite anatomical masks/images to this space (including those that were not automatically resampled in the `fmriprep` workflow).

We restrict our analysis to gray-matter voxels (GM), retaining only voxels with GM probability (calculated in fmriprep) greater than 50% to minimise partial-volume effects. We also exclude voxels with signal intensities below 70% of the mean fMRI signal intensity to avoid contamination by voxels with low signal caused by susceptibility and partial-volume effects.

ICA-based Automatic Removal Of Motion Artifacts (AROMA) was used to generate noise regressors, which were removed from the data using the non-aggressive variant of the method (56). Regressors were calculated on the spatially smoothed minimally preprocessed images (as described within `fmriprep` as a 6 mm FHWM kernel) and then applied to the unsmoothed preprocessed images.

Following ICA-AROMA, we extracted mean time courses from eroded masks (using a 3 *×* 3 *×* 3 erosion kernel) of the WM and CSF. The masks were generated by following Parkes et al. (41) and Power et al. (42), where CSF and WM ROIs were created from tissue probability maps in `fmriprep`. We eroded the WM mask five times and the CSF mask once. Erosion is crucial to avoid partial-volume effects from gray matter, which inflates the correlation between WM/CSF estimates and the global-mean signal (41, 57). We extracted these signals from the AROMA-denoised data, as performed in Pruim et al. (56).

The above steps are fairly standard and accepted de-noising techniques, but they are insufficient for removing widespread WSDs that are often tied to physiological and motion-related noise (40–43). To explicitly address the problem of WSDs, we implemented two different types of correction: Global signal regression (GSR), and Diffuse Cluster Estimation and Regression (DiCER), both of which have been shown to mitigate the influences of WSDs in different ways (43).

For GSR, we use the mean gray-matter signal as an estimate of the global signal, since the two are highly correlated and grey matter contributes the most to the global signal (42, 48). We thus refer to regression of this signal grey matter regression (GMR) for clarity. This method is by far the most commonly used to address WSDs in fMRI data.

Our second processing stream removed WSDs using DiCER. As shown in (43), GMR often results in incomplete removal of WSDs when the data are inspected under an appropriate re-ordering of the conventional carpet plot. DiCER iteratively removes WSDs by extracting representative signals for diffuse, weakly correlated clusters of voxels. The method shows superior denoising efficacy to GMR and can improve statistical sensitivity within datasets (43). Critically, it can be used to remove all WSDs, resulting in a “flattened” carpet plot. Whether this is desirable for fMRI processing remains debatable (43, 47, 58), but we use it here to evaluate model performance in data with minimal WSDs. Up to five noise regressors were estimated for each individual dataset with DiCER, as described in Aquino et al. (43).

We compare data processed with GMR and DiCER to data in which neither step has been applied, resulting in three parallel processing streams: (i) lenient– regression with the WM and CSF physiological signals, denoted as ‘+2P’; and (ii) moderate – regression with WM, CSF and GM signals, denoted as ‘+2P+GMR’. (iii) aggressive – regression with WM, CSF and DiCER regressors, denoted as ‘+2P+DiCER’. The first two models were applied after ICA-AROMA denoising in a single step using ordinary least squares regression implemented in `fsl_regfilt` and the last model was applied post (i), as DiCER targets residual WSDs in the data.

The data, including the minimally preprocessed data, were then detrended with a 2nd order polynomial and high-pass filtered at 0.005 Hz using AFNI’s `3dTproject`. This procedure resulted in four datasets for each subject, labeled ‘pre-ICA-AROMA’,’ICA-AROMA+2P’, ‘ICA-AROMA+2P+GMR’, ‘ICA-AROMA+2P+DiCER’.

#### Node definition and functional connectivity matrices

We defined brain regions using the the Desikan Killiany atlas (59), mapped on to cortical surface models of individual participants generated using Freesurfer (60). This cortical parcellaion was combined with the `aseg` segmentation of subcortical structures also provided by Freesurfer, resulting in 82 regions. The mean fMRI time series from each region was extracted from each of the four processing streams. Correlations between time series for each pair of regions were then estimated, resulting in a set of four connectivity FC matrices of size 82 *×* 82 for each subject. We note that the general points we make here will apply regardless of the specific parcellation chosen.

#### Structural connectivity matrix (SC)

We mapped structural connectivity using Human Connectome Project (HCP) data for 973 participants. Data were acquired on a customised Siemens 3T “Connectome Skyra” scanner at Washington University in St Louis, Missouri, USA using a multi-shell protocol for the DWI with the following parameters: 1.25 mm^3^ voxel size, TR = 5520 ms, TE = 89.5 ms, FOV of 210 *×* 180 mm, 270 directions with *b* = 1000, 2000, 3000 *s/*mm^2^ (90 per *b* value), and 18 *b* = 0 volumes. Structural *T*_1_-weighted data were acquired with 0.7 mm^3^ voxels, TR = 2400 ms, TE= 2.14 ms, FOV of 224 *×* 224 mm (61, 62).

The HCP data were processed according to the HCP minimal preprocessing pipeline, which included normalisation of mean *b* = 0 images across diffusion acquisitions, and correction for EPI susceptibility and signal outliers, eddy-current-induced distortions, slice dropouts, gradient nonlinearities and subject motion. *T*_1_-weighted data were corrected for gradient and readout distortions prior to being processed with Freesurfer (62). Tractography was conducted using the Fibre Orientation Distributions (iFOD2) algorithm, as implemented in MRtrix3 (63), which utilises Fibre Orientation Distributions (FODs) estimated for each voxel using Constrained Spherical Deconvolution (64–66). This approach can improve the reconstruction of tracts in highly curved and crossing fibre regions (64, 66).

Streamline seeds were preferentially selected from areas where streamline density was under-estimated with respect to fibre density estimates from the diffusion model (14). We used Anatomically Constrained Tractography to further improve the biological accuracy of streamlines (67). To create a structural connectivity matrix, streamlines were assigned to each of the closest regions in the parcellation within a 5 mm radius of the streamline endpoints (68), yielding an undirected 82 *×* 82 connectivity matrix per subject. The resulting matrices were averaged to form a group-average SC matrix which was normalised to have maximum connectivity 0.2.

#### Quality control methods

We are interested in understanding how DNMs perform following the application of different denoising (i.e. the removal of signals associated to non-neuronal fluctuations) pipelines to fMRI data. Implicit in this comparison is the notion that some pipelines may be more effective than others in removing WSDs and other sources of noise (40, 41, 43). We thus use several indices of denoising efficacy, defined at both the individual and group level. We note that our analyses are restricted only to subjects that pass strict inclusion criteria, as described in Aquino et al. (43), which includes only participants with low-to-moderate head motion (Mean framewise displacement: FD*<* 0.2, only allowing 20% of frames to pass FD= 0.25, and excluding subjects that have 5 mm FD spikes).

At the individual level, we use carpet plots to evaluate denoising efficacy. We construct these plots for each participant by taking each fMRI time series and performing a *z*-score time series for each subject. In these plots, WSDs appear as large vertical bands; the plots can thus be used to visually inspect the degree to which a given preprocessing pipeline has removed these deflections. Reordering the rows of the carpet plot can emphasise distinct structures in the data. As defined in Aquino et al. (43), we use two types of ordering: (1) an ordering of voxels by their correlation to the global signal (GSO), which can be used to identify gradients of such a correlation across voxels; and (2) an ordering based on hierarchical clustering (CO), as implemented in https://github.com/BMHLab/DiCER, which can be used to reveal additional, more complex WSDs (see (43) for details).

We use CO for the data, where carpet plots are shown at the level of individual voxels, and GSO for the models, where carpetplots are shown at the level of parcellated regions. A large collection of over 500 subjects of these visualisations for the UCLA and other cohorts is available at: https://bmhlab.github.io/DiCER_results/. We also quantify the “flatness” of each individual’s carpet plot using the variance explained by the first principal component (PC) of the voxel *×* time rsfMRI data matrix, denoted as ‘VE1’. High values of VE1 indicate that a large proportion of fMRI variance can be captured by a single component (PC1), consistent with the presence of dominant WSDs.

At the group level, we consider two measures of denoising efficacy. The first is the Quality Control–Functional Connectivity correlation (QC–FC), which is a commonly used benchmark (40–42) and is estimated as the cross-participant correlation between FC and mean FD at each edge in an FC network. The QC–FC correlation quantifies the association between inter-individual variance in FC and gross head motion, indexed by mean framewise Displacement (mFD) (57). An efficient denoising method will result in data that is less corrupted by head motion and thus lower QC–FC scores. As done previously (40, 41, 43), we estimate the QC–FC for each network edge, and summarise the findings by counting the percentage of *p <* 0.05 (uncorrected) QC–FC correlations.

## Modelling Methods

We evaluate the performance of three DNMs that are used widely and have been previously shown to reproduce empirical rsfMRI FC patterns (24, 26, 30, 31, 34–37, 51, 52, 69). These models are a subset of all available neural models but are representative of the classes of models used to fit empirical rsfMRI in the literature. The models have the following general form:

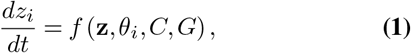

where neural activity *z*_*i*_(*t*) is described at node *i*, owing to a dynamical model *f* that depends on all other nodes (and populations) **z**, with parameters *θ*_*i*_ and structural connectivity matrix *C* scaled by global parameter *G*. Below we describe the details of each model.

### The balanced excitation-inhibition model

The balanced exhibition-inhibition (**BEI**) model uses a mean-field approach to simulate the collective behaviour of excitatory cell populations that are in “balance” with inhibitory populations (26). This firing rate model is an extension of the Wong and Wang (70) model that additionally tunes the excitatory-inhibitory ratio of each population such that each node in the system has a mean firing rate of 3 Hz, as suggested by invasive neurophysiological recordings (see (70) and (26)). This balance of excitatory to inhibitory connections ensures stability of the neural populations. The model comprises coupled stochastic differential equations that simulate the firing rates of excitatory (E) and inhibitory populations (I):

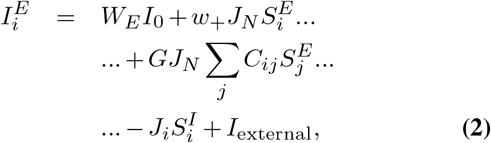

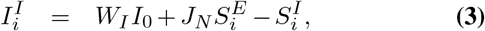

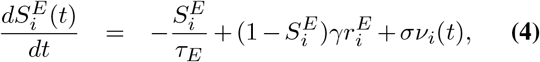

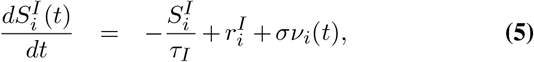

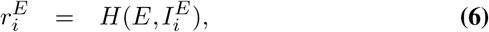

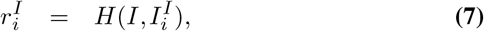

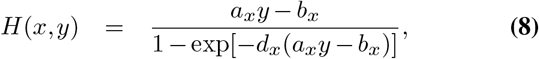

where the variable 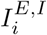 indicates ionic current, 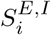 denotes the average synaptic gating variable, and 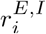 denotes the population firing rate. All these dynamic variables are described at excitatory and inhibitory populations (*E, I*) at node *i*. The parameters *W*_*E*_, *W*_*I*_ describe the overall scaling of the excitatory and inhibitory currents respectively, *w*_+_ describes the local excitatory recurrence, *J*_*N*_ is the excitatory synaptic coupling, *τ*_*E,I*_ are time constants for excitatory and inhibitory populations, *γ* is a kinetic rate constant. The function *H*(*x, y*) is the neuronal input-output response function for population *x* due to input current *y*, which is parametrised for population *x* with parameters *a*_*x*_, and *b*_*x*_. The function *H*(*x, y*) defines how the net current *I* induces a population level firing rate *r*.

The model simulates resting state activity as the neural response to a noisy input *σν*_*i*_(*t*), where *σ* scales the level of noise. We evaluated the model numerically using MATLAB with an Euler-Maryuami integration scheme with time step 0.01 ms. We note that the balancing of the dynamics – i.e., setting the local firing rate to 3 Hz – is achieved by adjusting the ratio of the influence of the inhibitory populations parametrised by the parameter *J*_*i*_. The value of *J*_*i*_ is estimated via a greedy search algorithm that adjusts *J*_*i*_ so that each node achieves an excitatory frequency of 3 Hz as described by Deco et al. (26) where the initial estimate for the search is the fixed point of the dynamical system (see Appendix). The parameters, and their nominal values are in Table 1. The algorithmic implementation can be found at http://github.com/KevinAquino/modelling_comparisons/.

**Table 1.**
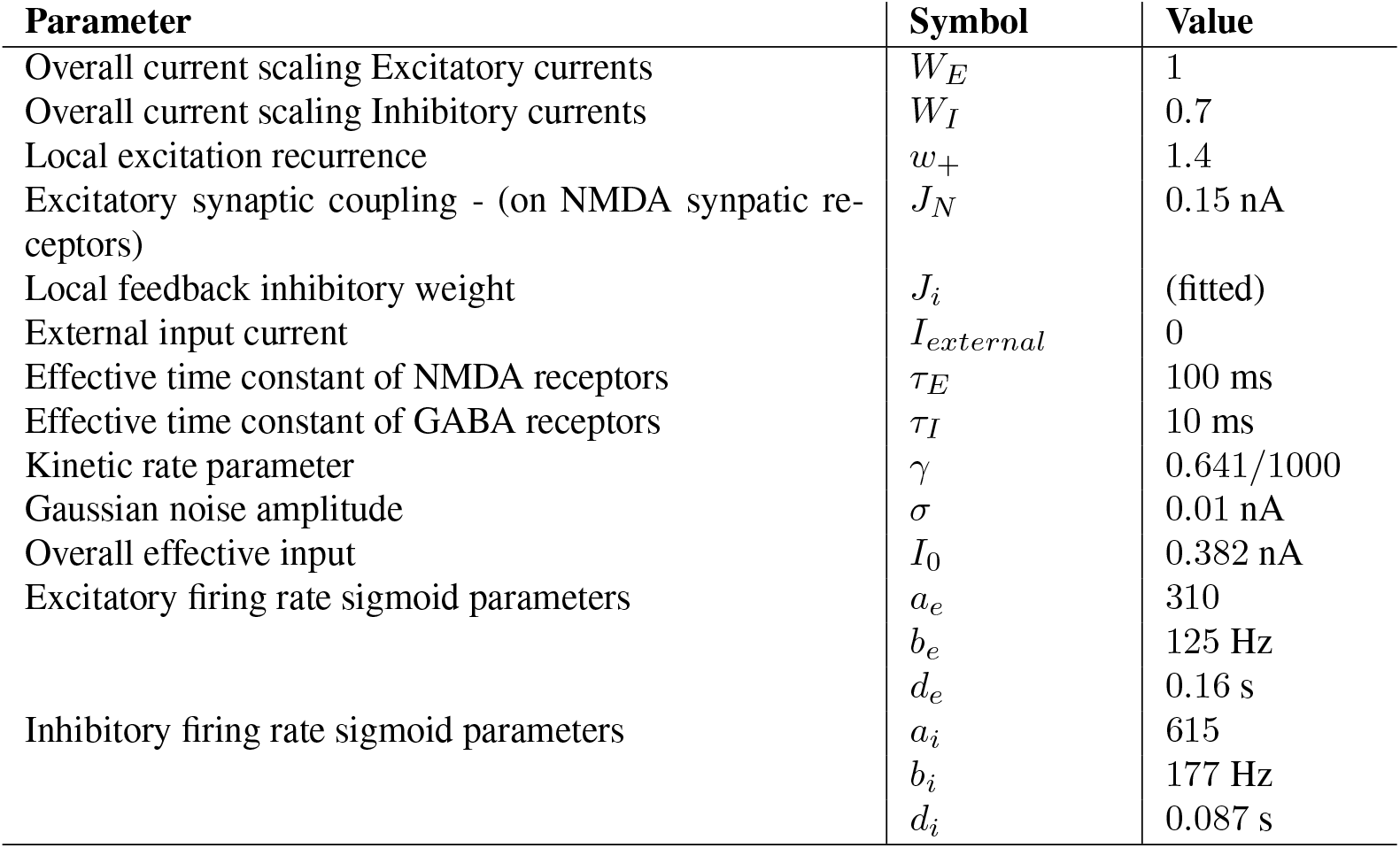
Parameters used for the Balanced excitation exhibition ratio model taken from (26).

To simulate BOLD dynamics, the output of the synaptic gating *S*_*E*_(*t*) is set as the neural drive,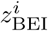, for the BOLD response (see below):

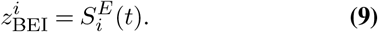

Finally, we note that *G* has no direct physiological analog and is fitted to the data so as to maximise the correlation between simulated and empirical FC estimates and minimise the discrepancy between predicted and empirical dynamic FC, as described below.

### Breakspear–Terry–Friston Model

The Breakspear Terry Friston (BTF) model simulates neuronal dynamics through a series of neuronal masses based on the Hodgkin-Huxley model (or equivalently the reduced Morris-Lecar model (71)) and has been used extensively in the literature (8, 24, 33, 35, 51, 52) including the influential study of the structure to function relationship of the human connectome(8). The main dynamical variables are the excitatory (*V*) and inhibitory (*Z*) membrane potentials, and the fraction of open sodium channels (*W*). At the level of node *i*, the dynamics are derived by using the equivalent electrical circuits of excitatory neurons, averaged over a small patch of cortex, that describe the conductance of sodium (Na), potassium (K) and calcium ions (Ca). The dynamical equations are

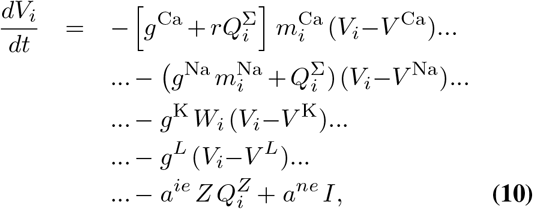

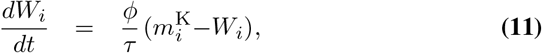

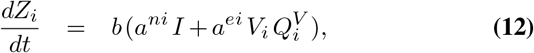

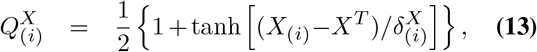

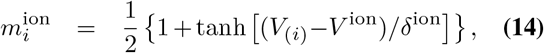

where the firing rate 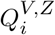 for excitatory and inhibitory populations are described via a sigmoid function that models the action potential at a population level, which is parametrised by the mean population thresholds *V* ^*T*^, *Z*^*T*^, and the variance of these thresholds *δ*^V,Z^. The parameter *g*^Na,Ca,K^ is the conductance of ion channels and *r* is the ratio of NMDA to AMPA receptors. The term 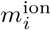 is the ratio of the open to closed ion channels that is described according to a sigmoid, parametrised by the mean Nerst potentials of the ion channels *V*^ion^ with the term *δ*^ion^ capturing the variance of these potentials.

We note that the ratio of open to closed ion Potassium channels is a special case and treated differently. As described in Breakspear et al. (33), this ratio varies dynamically to relax via an exponential decay in Eq. 11, where the decay parameter *τ* is the relaxation rate and *ϕ* is the temperature scaling factor. The parameters *a*^*ee*^, *a*^*ei*^, *a*^*ie*^, *a*^*ne*^, *a*^*ni*^ are the connection weights between inhibitory (*i*), excitatory (*e*), and external input (*n*) populations. The external population is modelled simply as a population that has input ionic current *I*. In this model, leaky currents are included that dissipate the excitatory membrane potential with appropriate parameters (of conductance and Nerst potential) with superscript *L*. The coupling between nodes (*G >* 0) is mediated through excitatory projections, where the net firing rate from the network onto a single node is given by

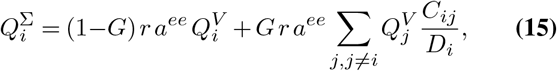

where *D*_*i*_ is the weighted degree of the node (*D*_*i*_ = ∑*j C*_*ij*_), and 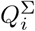 modulates the contributions of sodium and calcium excitatory membrane potentials. These equations form a nonlinear dynamical system that, for realistic biophysical parameters and relatively weak coupling (low *G*), exhibits chaotic dynamics with weakly synchronised activity (72). It is this regime that is typically used to model neural dynamics (8, 35, 51, 52). As in the BEI model, *G* has no direct physiological analog and is chosen to maximise similarity between simulations and data (see below for more details).

The BTF model simulates spontaneous activity as the cortical response due to unstructured white noise *a*^*ne*^*I*, where *I* is the nonspecific subcortical current. As the system of equations exhibit emergent chaotic behaviour, it is only necessary to include random initial conditions across all nodes to reproduce ongoing dynamics as opposed to the constant drive of noise *σν*_*i*_(*t*) in the BEI model (and the Hopf model below) where the dynamics relax back down to the fixed point following excitation. Specifically, at each run of the BTF model, we set initial random (with a uniform distribution) values for *V, W* and *Z* at *t* = 0 that range between: [*−* 0.6, 0.2], [0, 0.6] and [*−*0.03, 0.13] mV respectively.

The model parameter values are listed in Table 2. The model was numerically integrated using a 4th order Runge-Kutta scheme within the Brain Dynamics Toolbox (73) in MATLAB (details within the provided toolbox).

**Table 2.** Parameters for the BTF model, using the values from Honey et al. (35).

| Parameter | Symbol | Value |
| --- | --- | --- |
| Ionic gains (L is the leaky current) | $g^{Ca}, g^K, g^{Na}, g^L$ | 1.1, 2, 6.7, 0.5 |
| Mean Nerst potentials | $V^{Ca}, V^K, V^{Na}, V^L$ | 1, -0.75, 0.53, -0.5 |
| Variance of Nerst potentials | $\delta^{Ca}, \delta^K, \delta^{Na}$ | 0.15, 0.30, 0.15 |
| Mean excitatory population voltage threshold | $V^T$ | 0 |
| Mean inhibitory population voltage threshold | $Z^T$ | 0 |
| Variance of excitatory population voltage threshold | $\delta^V$ | 0.65 |
| Variance of inhibitory population voltage threshold | $\delta^Z$ | 0.65 |
| Ratio of NMDA to AMPA receptors | $r$ | 0.25 |
| Temperature scaling factor | $\phi$ | 0.7 |
| Decay time constant of Potassium channel dynamics | $\tau$ | 1 |
| Decay time constant of Inhibitory populations | $b$ | 0.1 |
| Nonspecific subcortical input current | $I$ | 0.3 |
| Connection weights between neural populations and subcortical inputs | $a^{ee}, a^{ei}, a^{ie}, a^{ne}, a^{ni}$ | 0.36, 2, 2, 1, 0.4 |

To derive BOLD dynamics, the absolute rate of change of the excitatory membrane potential is used as a proxy for the glutamate uptake that drives the neurovascular response, i.e.

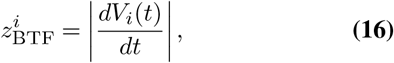

where the drive 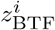 is used in a BOLD forward model (see below).

### The Hopf bifurcation model

The BEI and BTF models attempt to simulate neuronal activity as the evolution of a set of explicitly parametrised biophysical processes. The BEI models synaptic dynamics driven by noise whereas the BTF models ion channel dynamics. The BEI model exhibits stochastic noisy dynamics without explicit oscillations and the BTF model exhibits oscillatory chaotic dynamics. An alternative approach is to model oscillations, noise, and their combination; i.e., noisy oscillations. This can be achieved using the supercritical Hopf bifurcation oscillator - which is a normal form to model the dynamic behaviour present in many bio-physically realistic models (34, 74–78). The explicit model described in Deco et al. (34) describes dynamics for two variables *x*_*i*_, *y*_*i*_ per node *i* through a series of coupled stochastic differential equations:

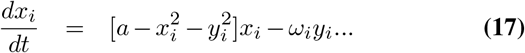

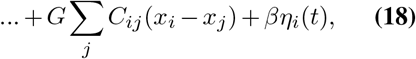

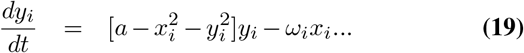

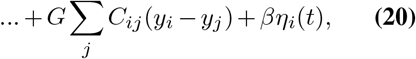

where *a* is the bifurcation parameter that, at the node level, tunes the system to be in one of three regimes. At the individual node level without coupling (*G* = 0), if *a >* 0 the system is in an oscillatory regime with frequency *ω*_*i*_, and if *a <*= 0 the system lies around a fixed point perturbed by Gaussian noise *η*(*t*) (with standard deviation *β*). When coupling between nodes is introduced (*G >* 0) through the term *G ∑j C*_*ij*_(*y*_*i*_ *− y*_*j*_), the dynamics of the system changes its characteristics in accordance with the bifurcation parameter *a*. When *a <* 0, the dynamical system exhibits noisy deviations about a stable fixed point, when *a ≈* 0 the system exhibits noisy oscillations, and when *a >* 0 the system exhibits oscillatory behaviour. The frequency *ω*_*i*_ is estimated from resting state data itself by calculating the peak frequency for all nodes within the narrow frequency band from 0.04 to 0.07 Hz and averaged over all subjects.

This model requires two parameters to be fitted: the global coupling parameter *G*, and the bifurcation parameter *a*. We use *a* = *−* 0.001, as per prior work (34) and fitted *G* to maximise similarity between simulations and data (see below for more details for all homogeneous models). The last parameter *β* determines the strength of the noise term which is set at 0.02. The dynamical equations are solved with MATLAB, using an Euler Muryami integration scheme (with time step 0.01 ms with the code available at http://github.com/KevinAquino/modelling_comparisons/ with the parameters described in Table 3.

**Table 3.** Parameters for the Hopf model, using the values from Deco et al. (34).

| Parameter | Symbol | Value |
| --- | --- | --- |
| Standard deviation for Gaussian noise | $\beta$ | 0.02 |
| Inherent frequency at each node $i$ | $\omega_i$ | Estimated directly from data |
| Global bifurcation parameter | $a$ | -0.01 (homogenous case) |

**Table 4.** Parameters for the Baloon model derived from Deco et al. (26).

| Parameter | Symbol | Value |
| --- | --- | --- |
| Signal decay rate | $\kappa$ | $0.65 \text{ s}^{-1}$ |
| Flow elimination constant | $\gamma$ | $0.41 \text{ s}^{-2}$ |
| Mean transit time | $\tau$ | 0.98 |
| Grubb's exponent | $\alpha$ | 0.32 |
| Resting oxygen extraction fraction | $E_0$ | 0.34 |
| Resting blood volume | $V_0$ | 0.02 |
| BOLD parameters | $k_1$ | $7E_0$ |
| | $k_2$ | 2 |
| | $k_3$ | $2E_0 - 0.2$ |

Since the oscillations are set to the frequency of the BOLD response, there is no need for a BOLD forward model with the Hopf model(34).

### Imposing Heterogeneity

The presented models impose uniform node parameters. Such models provide moderate predictive power for static FC but can have limited efficacy in capturing dynamic (i.e. time-varying) properties of FC (34). To improve model fitting to both static and dynamic aspects of FC, we can increase the degrees of freedom of a given model by introducing heterogeneity to the node-level parameters ((36, 38)). This is because heterogeneity of node parameters can cause changes in local dynamics (e.g., temporal autocorrelation, excited frequencies per node, and the ratio between noise and oscillations), which can disrupt globally synchronous activity and expand the dynamic range of the system.

To limit computational burden, we evaluate a simple form of heterogeneity within the Hopf model by varying the value of the bifurcation parameter *a* at the node level and specifying a node specific bifurcation parameter *a*_*i*_, yielding:

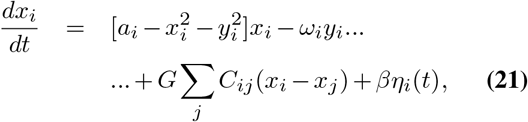

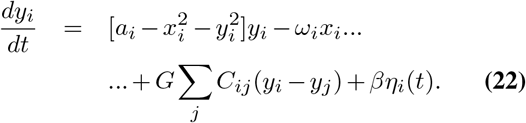

The model is estimated following Deco et al. (34), where for each *G* we optimise *a*_*i*_ using a greedy search algorithm to optimise the model fit to dynamic aspects of FC (34) (see below) over *N* parameters.

Another way to introduce additional degrees of freedom is by adjusting the connection weights of the structural connectivity matrix *C*_*ij*_, a concept we denote as “edge heterogeneity”. As detailed in Gilson et al. (69), the aim is to modify the existing connectivity *C*_*ij*_ between two nodes *i* and *j* to arrive at an effective connectivity *E*_*ij*_ which improves the match of the dynamics between the simulated and the recorded dynamics. We introduce this edge heterogeneity to the node-homogeneous Hopf model, using the optimisation procedure described below. We denote as Hopf-node and Hopf-edge and as the node-heterogeneous and edge-heterogeneous Hopf models, respectively.

### BOLD forward model

The neural dynamics resulting from both the BTF (Eq. 16) and BEI models (Eq. 1) are translated to BOLD activity via the Balloon-Windkessel model (27, 28, 79), using (Eq. 16) and (Eq. 1) as neuronal drivers to the system (See appendix). Although the balloon model is not a physiologically faithful model of vascular hemodynamics (80–83), it can reproduce the nonlinear BOLD dynamics in task fMRI and is routinely used and fitted in dynamic causal modelling (27). We also use this model in place of more realisitic models of the hemodynamic response(83–85) for adequate comparisons with previous studies of large-scale brain modelling.

As the Hopf bifurcation model simulates general dynamics it does not need a forward model as the dynamics of *x*_*j*_ can be set at the time scale and properties of BOLD fMRI fluctuations.

For all models, a total of 5 minutes of spontaneous activity was simulated and downsampled to a sampling rate of 2 s, mirroring the time scale of typical single-band resting state fMRI data and the specific dataset considered here.

### Parameter Optimization

While all of the presented models have parameters that are based on physiological recordings (**BTF, BEI**) or direct measurements of dynamics (**Hopf**), they additionally require a specification of the global scaling hyperparameter *G*. This parameter has no direct analogue with data and is chosen to optimise the match between simulated data and experiment. Below we describe the metrics that we used to quantify model fit and the general approach for optimising *G* in homogeneous models (models with uniform node parameters). We then describe further optimisation steps taken for the Hopf-node and Hopf-edge heterogeneous models. For each of the models considered, at each possible value of *G*, the resting state simulation is run for 304 seconds (post initial transients in the dynamic models) and run 108 times to match the number of subjects in the empirical data (that pass quality control measures).

#### Assessing model performance

We evaluate model performance with respect to static and dynamic FC properties as has been done previously (26, 34). For static FC, we first estimate Pearson correlations between every pair of simulated regional time series in the model to generate a synthetic FC matrix. We then quantify static FC model fit as the correlation, *R*_*s*_, between the upper triangles of the z-transformed synthetic (FC_s_) and empirical (FC_e_) FC matrices:

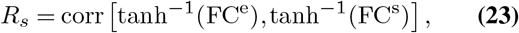

as performed in Deco et al. (26). This metric ranges from *−* 1 to 1, where *R*_*s*_ = 0 implies the model and data are completely uncorrelated an |*R*_*s*_|= 1 implies an exact correlation of the functional connectivity structure between data and model.

To fit dynamic properties of FC, we focus on the phase-derived functional connectivity dynamics (FCD). This measure is an improvement on previous FCD metrics (34) as it is not based on amplitude and evaluates phase-lagged coupling structure over time. First, the time series at each node is filtered with a 2nd order band-pass Butterworth filter (between 0.04 and 0.7 Hz). The instantaneous analytic phase *θ*_*j*_(*t*) from each node *j* is estimated by using the Hilbert transform and the cosine of phase difference between node *i* and *j*, Δ_*c*_(*i, j, t*) is then calculated as follows:

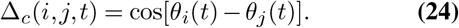

This metric is the instantaneous phase difference between node *i* and *j* at time *t*, and thus captures the level of synchrony between nodes at each time point.

To determine the time-varying aspects of this synchrony across all nodes, we compute the dot product (*ϕ*_*uv*_) between the upper triangle of the matrix in Eq. 24 (i.e. across all nodes) in one time instance *τ*_*u*_ vs another time instance *τ*_*v*_ i.e.

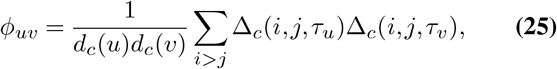

where:

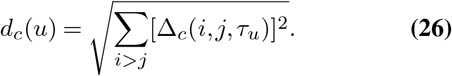

We can interpret *ϕ*_*uv*_ as the similarity of the global synchrony between time *τ*_*u*_ and time *τ*_*v*_.

The *ϕ*_*uv*_ metric will exhibit block structure off the main diagonal if the overall network dynamics, as revealed by the pairwise phase differences between nodes, are similar between a given pair of time points. The distribution of *ϕ*_*uv*_ (i.e. across all time lags) thus characterises general fluctuations of the system. As the distribution of *ϕ*_*uv*_ in a single instance of either a model simulation or the data depends on the specific synchrony transitions, the distribution of *ϕ*_*uv*_ is analysed over multiple realisations of the model or in data it is aggregated across all subjects at a group level.

We quantify the similarity of fluctuations between model and data at the level of *ϕ*_*uv*_ distributions using the Kolmogorov-Smirnoff (KS) statistic - a measure of the “distance” between two distributions. A low score on this statistic indicates more similar fluctuations. We denote the KS statistic metric as the functional connectivity dynamics - FCD.

#### Optimising homogeneous models

In the homogeneous models specified here, we only require the tuning of the hyperparameter *G*. This parameter is tuned to maximise *R*_*s*_ (fit to static FC) and minimise FCD (fit to dynamic FC). This one-parameter optimisation has been used previously in Deco et al. (34) and in that study *R*_*s*_ typically monotonically increases (or plateaus) whereas FCD is minimised. The use of two metrics allows a tight constraint: to maximise *R*_*s*_ and minimise FCD.

#### Optimising the node-heterogeneous model

As described above, we impose heterogeneity at the node level by varying the local bifurcation parameter *a*_*i*_ of the Hopf model, as described in Equations 21 and 22. To find values of *a*_*i*_ we use the approach in Deco et al. (34) where, for a given coupling *G*, the parameter *a*_*i*_ for node *i* is varied to match the observed local power spectra – specifically to match the ratio of low frequency power relative to high frequency power. The rationale is that high frequencies are related to noisy processes (as the BOLD response is mostly a low-pass filter) compared to low frequency oscillations, which are more strongly determined by network activity. This optimisation is achieved using a gradient descent algorithm, where we vary *a*_*i*_ as

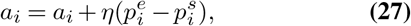

where *η* = 0.1, and both 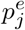 and 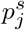 are the ratios of the totalpower in the narrow band between 0.04 and 0.07 Hz and the high frequency power 0.07 and 0.25 Hz for the empirical and simulated data respectively. Note that the empirical ratio 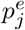 has been averaged over all subjects. This optimisation drives each node toward the oscillatory regime if 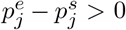 or toward the noise regime if 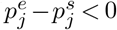. The stopping criterion is 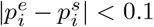. Once the model parameters have been optimised for a particular *G*, the model is then run as above with these optimised parameters (i.e, run 108 times with each run lasting 304s to match the data).

#### Optimising the edge-heterogeneous model

To introduce heterogeneity at the level of edges (i.e. variation of connectivity weights) we vary *C*_*ij*_ in the Hopf model as follows. First, the connectivity strengths that have non-zero values in *C* are used as a mask such that only those edges will be modified, with the model of effective connectivity i nitialised as *E*_*ij*_ = *C*_*ij*_. Second, for each value of *G*, the model is estimated. Third, after each model simulation the edges *E*_*ij*_ are adjusted such that

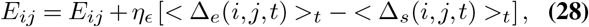

where the strength of edge variation *η* _*ϵ*_ = 0.01, and where *<* Δ_*e*_(*i, j, t*) *>*_*t*_ and *<* Δ_*s*_(*i, j, t*) *>*_*t*_ are the time-averaged phase cosine difference (as per 24) between node *i* and node *j* for the experimental data, *e*, and simulated data, *s*, respectively. This procedure forces pairs of nodes to match the level of synchrony exhibited in the data and is repeated for all edges in the mask, constituting one iteration of the optimisation. This adjustment is repeated until the root mean square of *<* Δ_*e*_(*i, j, t*) *>*_*t*_ *−<* Δ_*s*_(*i, j, t*) *>*_*t*_ falls below 0.01, thus optimising the match between model and data with respect to inter-regional synchrony. If the edge adjustment yields an edge to have *E*_*ij*_ *<* 0, then this edge is set to 0. This greedy optimisation converges and is a rough approximation of the procedure in Gilson et al. (69), however here *E*_*ij*_ is symmetric.

As the parameters for *E*_*ij*_ span all possible edges, i.e. (*N × N − N*)*/*2 parameters, this procedure can be prone to over-fitting (as is the case for the node level heterogeneity but we exclude that comparison in this study). We thus fitted *E*_*ij*_ on 80% of the sample and tested model performance on the remaining 20% of the data. This cross-validation step is repeated 108 times, i.e., the 80% random splitting, where the subject selection is without replacement in the test–retest split.

## Results

The results are structured as follow. First, we investigate the spatiotemporal characteristics of fMRI data and properties of data denoising pipelines that can influence model fi ts. Second, we characterise the spatiotemporal characteristics of the homogeneous DNMs and compare the gross features of the models with the data. Third, we quantitatively evaluate homogeneous model fits to the data in the static and dynamic regimes, where the data have been preprocessed using different denoising algorithms. Finally, we evaluate the performance of heterogeneous models in capturing aspects of the data that are not accurately reproduced by homogeneous models.

### Spatiotemporal properties of rsfMRI data

To enable us to better understand model performance we present the data that is the target of model simulations. Specifically, we show here how different preprocessing procedures can affect the spatiotemporal structure of fMRI data and that will thus lead to different models. Figure 2C, shows carpetplots for a “high-motion” subject from the UCLA cohort (sub-10274) following basic preprocessing (pre-ICA-AROMA), along with corresponding global signal and motion traces (Figs. 2A and B, respectively). This individual shows pronounced WSDs that are at times tied to motion events as captured by the mean framewise displacement (FD) (red rectangles). Additional WSDs that are not linked to motion (blue rectangles) can be linked to physiological events such as respiration (44, 49) or heart-rate variability (86). A robust de-noising method – ICA-AROMA with regression of white and CSF signals (+2P) — can reduce motion-related contamination but often leaves significant residual WSDs (44, 87), as shown in Fig. 2D. The effect of these WSDs on network architecture is shown in the FC matrix in Fig. 2G, where globally positive correlations are evident. Application of GMR, as seen in Fig 2E, can help mitigate the effects of WSDs, but centres the distribution of FC values on zero (88, 89) and thus introduces negative correlations throughout, some of which may be spurious (89, 90) (see Fig. 2H and Fig. 3H). Moreover, GMR will only be effective in removing WSDs if the signal deflections are in phase (especially if they are *π* radians out of phase). In some individuals, WSDs can have a biphasic (in this dataset they are out of phase by *π* rad.) appearance, with some voxels showing increased signal and others showing decreased signal. In these cases, GMR will be ineffective (43), as shown in Fig. 3 in blue where at times the WSDs are expressed in two clear phases - i.e., the top half has the opposite sign of the bottom half, and GMR does not have a significant impact on WSD structure at these periods (compare the residual WSD structure in Fig. 3D vs Fig. 3E in the blue vertical stripes).

DiCER was developed to identify and remove these more complex WSDs that are not adequately removed by GMR (43). On application of DiCER to the ICA-AROMA+2P output from all participants, the carpet plots in Figures 2F and 3F show the effective removal of WSDs, whilst revealing FC matrices in Figs. 2 I and 3 I that have structure without zero-centring the FC matrix by construction.

Figure 4 further shows the differences between preprocessing strategies with respect to group-level quality metrics. First, we can see that the correlation structure of the sample mean FC matrices differ substantially as a function of denoising strategy. Following ICA-AROMA+2P, the data is globally positively correlated (Fig. 4A). ICA-AROMA+GSR reveals more structure, with an approximately equal proportion of positive and negative correlations (Fig. 4B). Finally, ICA-AROMA+DiCER also shows reduced global correlations, with similar positive FC structure to GMR but reduced magnitude of negative FC estimates (Fig. 4C).

**Fig. 4.**
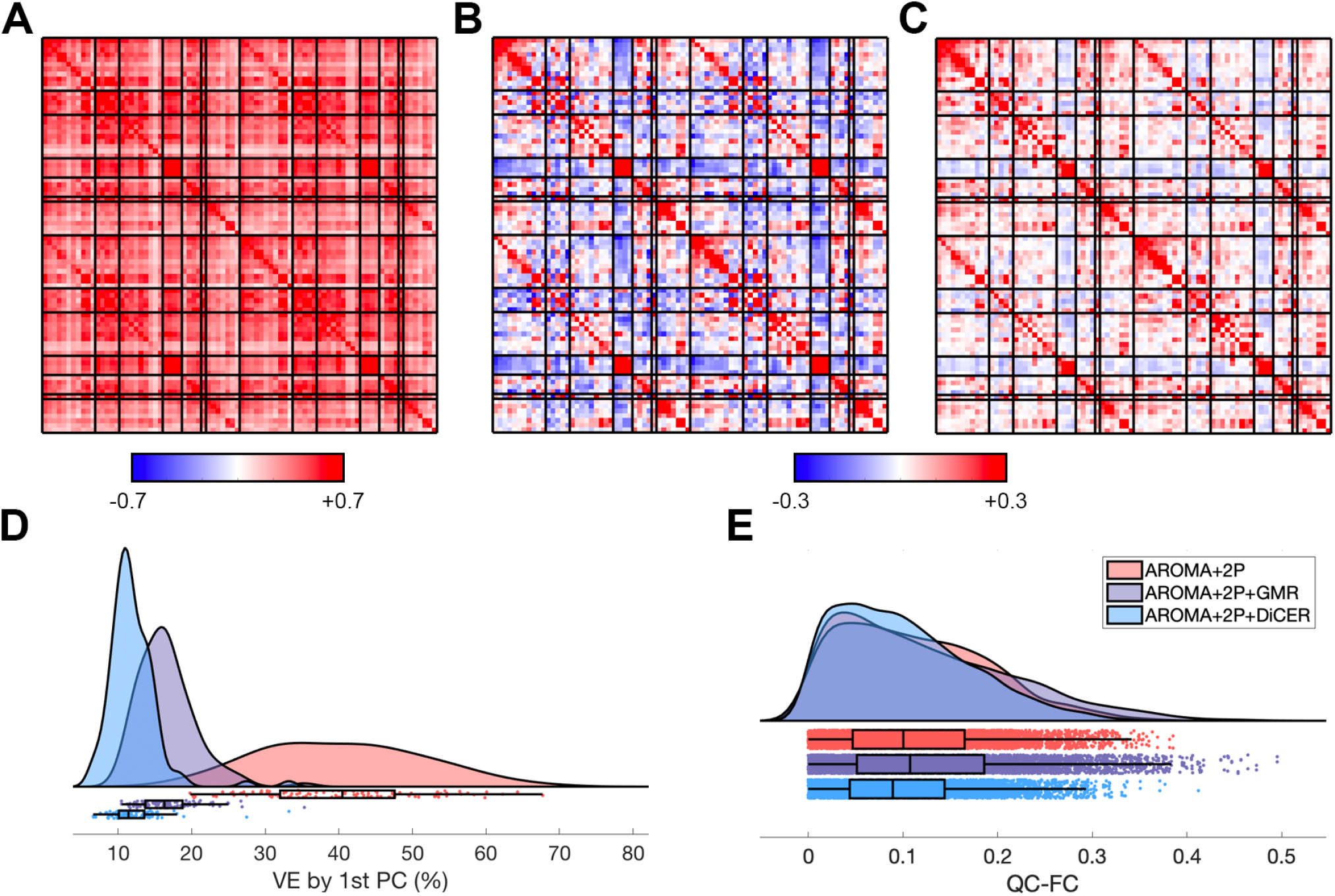
Group-level quality control metrics applied to the resting-state fMRI data. Panels **A, B**, and **C** show the sample mean FC matrices of the subjects processed under ICA-AROMA+2P,ICA-AROMA+2P+GSR, and ICA-AROMA+2P+DiCER respectively. Note that the colourscale changes as the range of FC values reduces for the last two pipelines. Panel **D** shows the distribution VE1 across individuals, and **E** shows the distribution of QC-FC estimates across all subjects with the three denoising pipelines. In both D and E, the distributions are represented as raincloud plots where the top curves are kernel density estimates and the bottom shows the scatter with box plots overlaid (91).

As shown in Figures 2 and 3, the three de-noising pipelines differ in the extent to which they remove WSDs from the data; i.e., they extent to which they “flatten” the carpetplot. One metric that can capture the flatness of the carpetplot is the proportion of variance explained by the first principal component of the data matrix, VE1 (see methods). Plotting VE1 for each individual, we see a clear difference as a function of data denoising (Fig. 4D), with VE1 being substantially higher for ICA-AROMA+2P, followed by ICA-AROMA+2P+GSR and lowest following ICA-AROMA+2P+DiCER. Average QC–FC estimates also tend to be lower following ICA-AROMA+2P+DiCER (Fig. 4E).

For present purposes, it is sufficient to note that the application of these four pipelines yields a gradient defining the degree to which the empirical data are dominated by WSDs, with base preprocessing being the worst, followed by ICA-AROMA+2P, then ICA-AROMA+2P+GMR, and finally ICA-AROMA+DiCER.

### WSDs in dynamic neural models

As previously described in the methods, a key feature of all dynamical models presented here is the global coupling parameter *G*, which changes the dynamics of DNMs from uncoupled dynamical nodes (*G* = 0) through to globally synchronised activity (*G >* 4 in the models considered here). Given that WSDs influence the level of global correlation in the data, and that different preprocessing pipelines show varying efficacy in the removal of these WSDs (e.g., compare panels G and I in Figs. 2 and 3), we can thus conclude that any WSDs in the data will have a major influence on (1) the precise value of *G* that is chosen during model fitting; and (2) the resulting estimate of model fit. More specifically, we can expect that data with prominent WSDs will be optimally fit by models with high *G* and that model fits will be higher under such circumstances. To understand this behaviour in more detail, we now examine the spatiotemporal structure of synthetic signals generated under the BEI, BTF, and Hopf models in the same way experimental data in the previous section. Typical measures such as model fits are described later in the results. Figure 5 A shows carpet plots for the BEI model at different values of the global coupling parameter *G*. Note that we have not explicitly fitted the model to fMRI data; we just simulate BEI dynamics on an empirical SC (of size 82 *×* 82) for different levels of *G*. Visually, it is evident that the number and magnitude of WSDs increase as *G* increases, and this is reflected in the global structure in the FC matrices in Fig. 5 B. As shown in Fig. 5 C VE1, an index of global synchrony in the data and a proxy of the prominence of WSDs, increases as *G* increases. Figure 5 C also shows that if we aim to match VE1 of the model with empirical data (in this case, the mean VE1 across the sample of subjects), we need to choose different values of *G* for data processed with different pipelines. More specifically, and in line with our hypothesis, higher values of *G* are required to match the VE1 of empirical data processed with pipelines that leave greater residual WSDs. For instance, when the data are processed with ICA-AROMA+2P, the BEI model matches empirical VE1 at *G* = 3 whereas when the data are processed with ICA-AROMA+2P+DICER VE1 is matched at *G* = 1.5. This behaviour is also evident in the BTF model (Fig. 5 D–F) and the Hopf bifurcation model (Fig. 5 G–I). Note that the transition from uncoupled noisy nodes to a synchronous pattern increases gradually in the BEI model (Fig. 5 C) and in the Hopf bifurcation model (Fig. 5 I) and is more abrupt in the BTF model (Fig. 5 F), where the model shows globally coherent activity past *G ≈* 3.5. We note two pathological behaviours of high *G*: firstly, in the BEI model at *G* = 3.6 (Fig. 5 A) a transient ie evident near the start of the time series which arises when the simulation diverges 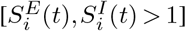, causing the membrane potential to reset 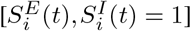 before the simulation continues on. Secondly, in the BTF for *G >* 3.6, the model can switch from weakly synchronized to fully synchronized as the model can take a long time to reach a synchronized limit cycle. Together, these two behaviours can cause some simulations in the BTF and BEI to not be fully synchronized which causes the variance of *V E*1 is higher and less predictable at high *G*.

**Fig. 5.**
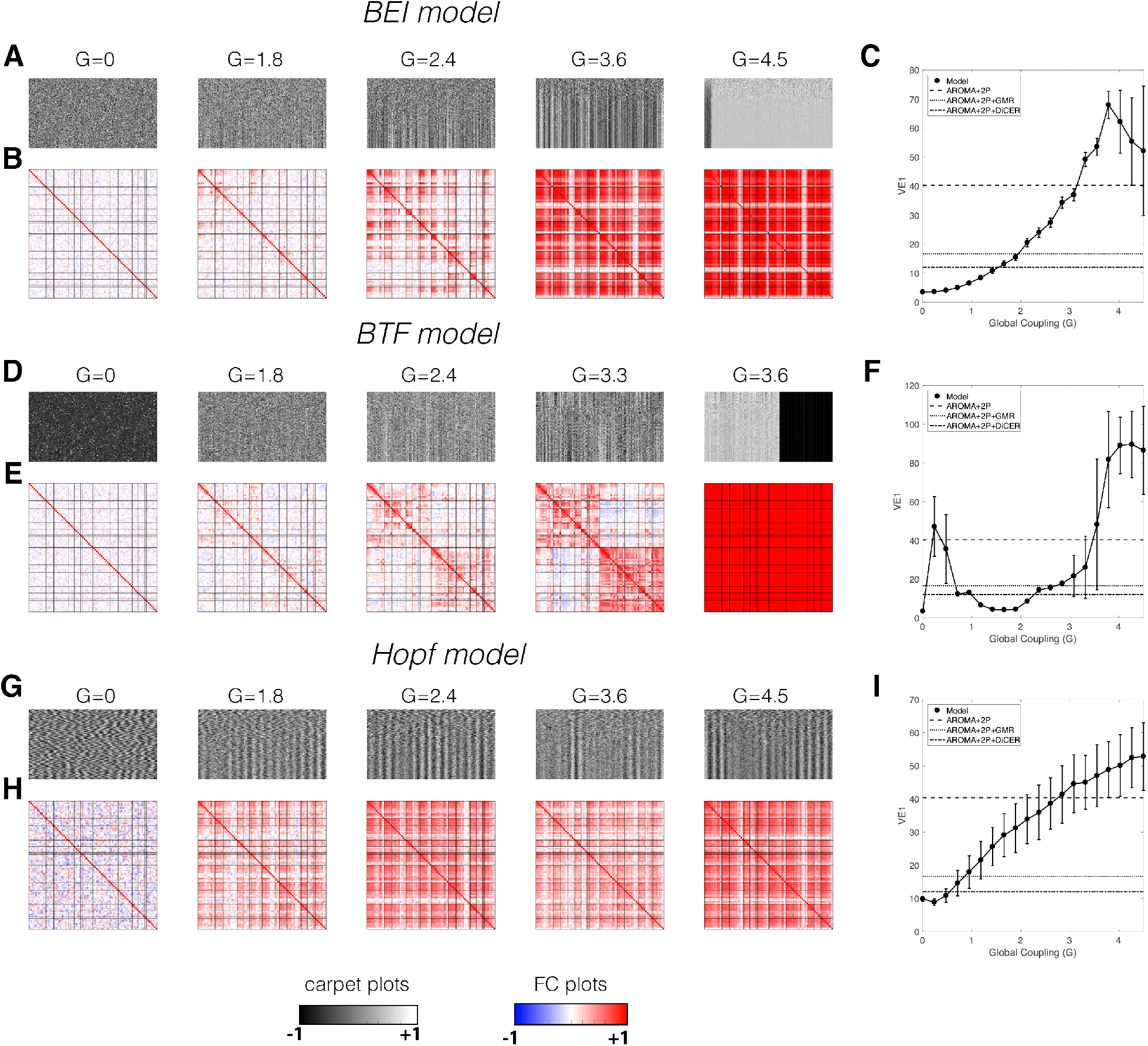
Widespread signal deflections in biophysical models of resting-state fMRI. In the three models presented here we show spatiotemporal carpet plots and FC matrices as a function of the global coupling parameter *G* in (A,B), (D,E) and (G,H) for one instance of the BEI, BTF, and Hopf bifurcation models, respectively. Quantitative metrics of variance explained by the first PC (VE1) are shown in (C), (F), and (I) for BEI, BTF, and Hopf bifurcation models, respectively. As described in the methods, the models were run 108 times and for 304s, mirroring the data. The curves for the model represent the mean metric values, with the error bars representing standard deviation at each *G*. The horizontal lines on the metrics (C,F,I) represent the mean VE1 of the empirical fMRI data under the three different denoising pipelines.

These results indicate that data with prominent WSDs require models to be biased toward the synchronised regime, whereas data with fewer WSDs push the model to the asynchronous regime (i.e. lower *G*). We stress that we have used exactly the same raw dataset for comparison. The only difference concerns how those data have been denoised.

### A noisy degree model of resting-state fMRI

The extent to which all WSDs in resting-state fMRI data represent physiological noise or actual neuronal dynamics in empirical data remains a topic of debate (43, 44, 48)). One interpretation of the results presented in Fig. 5 is that WSDs can arise as an intrinsic property of dynamics unfolding on a connectome, given a sufficient level of global coupling, since the models themselves contain no structured non-neuronal signals and WSDs are evident across all models, regardless of the underlying dynamics.

While this interpretation supports the validity and utility of whole-brain modelling, our findings raise the question of whether the models are only fitting globally coherent signals fluctuations. In other words, can a naive model of pure global synchronization fit the data just as well? Since data with large WSDs are routinely utilized in modelling, are models biased toward fitting globally synchronous activity?

To explore this issue further and better understand the dependence between WSDs in the data and model fits, we propose a simplified model. We start with the observation that, within the synchronized regime, DNMs show major, globally coherent WSDs, as shown by the large vertical stripes in Figs 5 A,D,H with *G* = 3.6, *G* = 2.4, 1.8 *≤G ≥*4.5 respectively. To construct a model of this behaviour, we first note that in all three DNMs considered here, the local dynamics at node *i* are modulated by the influences summed (or averaged) over neighbouring nodes *j* weighted by *C*_*ij*_ i.e., by a general factor:

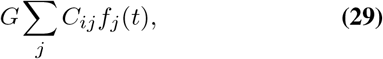

where *f*_*j*_(*t*) is a generic function that describes some output node *j* expresses to node *i*. In most models *f*_*j*_(*t*) depends on the dynamics of the excitatory population at node *j* projected to node *i*. Thus, if all nodes share a common signal *σ*(*t*), the dynamics at node *i* will be modulated by this common factor, a local noise term, and a regulatory term (expressing the natural decay of the signal). We can thus present a simple model that includes a common signal and a simple Gaussian noise process at each node, *N*_*i*_(0, 1) (owing to noisy inputs in all the three models). The model is expressed as:

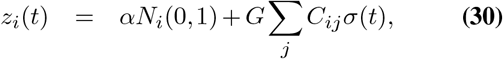

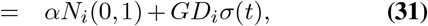

where *D*_*i*_ is the weighted node degree, and *α* determines the strength of the local noise term, which we set at *α* = 1*/*2 when considering z-scored common signals *σ*(*t*) (note that Eq. 31 should technically be derived with a rate of change of *z*_*i*_ and a self-regulation term, but we leave it in this form for simplicity).

This baseline model is derived through consideration of neural dynamics but we can substitute *z*_*i*_(*t*) with the BOLD signal *B*_*i*_(*t*) by setting the common *σ*(*t*) to have properties of typical BOLD global signals, which can be achieved by assuming a linear relationship between neural and BOLD signals. For each instance of the model, *σ*(*t*) is a random white signal for which we take a moving average of 10 seconds (5 volumes) to match the temporal autoocorrelation of the empirical data.

A single instance of this noisy degree model using *σ*(*t*) as the global signal from subject sub-10274 is shown in Fig 6.

**Fig. 6.**
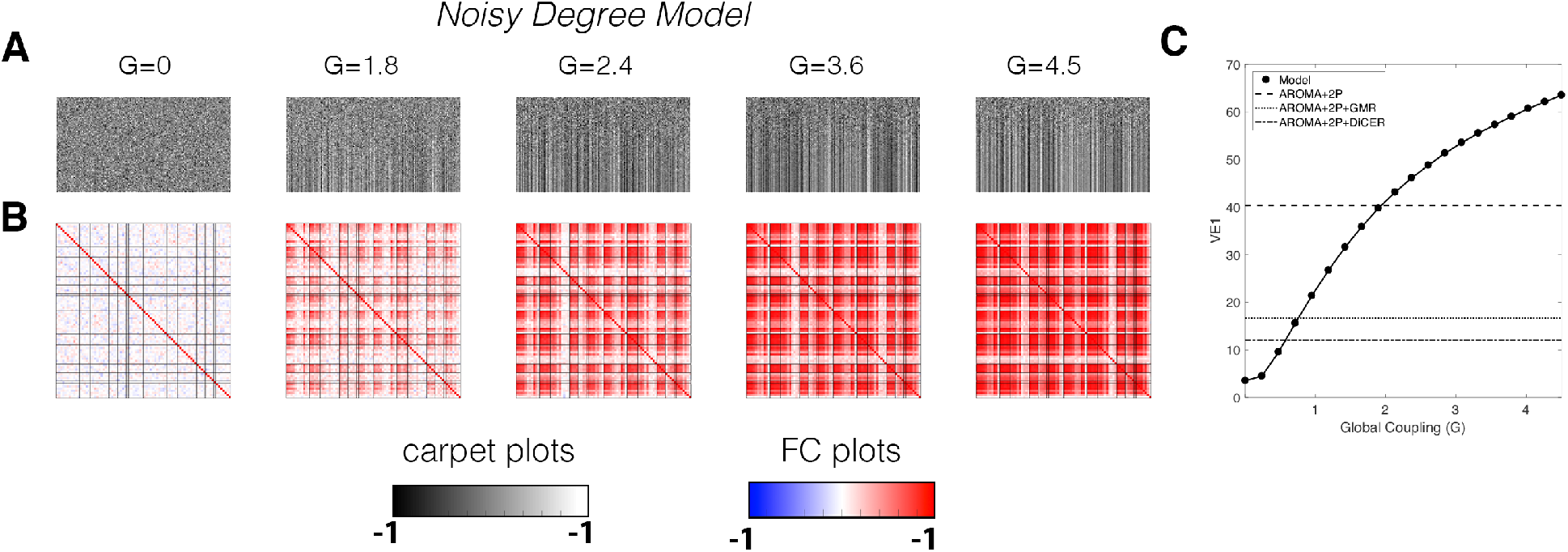
WSDs in the noisy degree model. (A) carpet plots and (B) functional connectivity matrices as a function of the global coupling parameter *G*. (C) Variance explained by the first PC (VE1), with solid line indicating the mean VE1 across 108 runs simulated at 304s with the error bars representing the standard deviation of these runs. Horizontal lines represent the mean VE1 under the three different denoising pipelines.

In the synchronized regime, the model generates carpet plots and FC matrices that are qualitatively similar to the BEI, BTF and Hopf models, and also the empirical data following ICA-AROMA+2P (cf. Fig. 2). Thus, a simple model that includes a global signal modulated by node degree shows dynamics that mimic typical DNMs in the synchronized regime, and the FC derived from this noisy degree model is virtually identical to the FC structure obtained with the BEI model (*r* = 0.99) at *G* = 3.6.

For all DNMs in the weakly synchronized regime (i.e., lower values of *G*), fewer WSDs are evident in the carpet plots, which is consistent with the empirical properties of data denoised with either GSR or DiCER.

### The influence of GMR on model dynamics

Since the DNMs we consider here explicitly simulate globally coherent neuronal fluctuations, a key question is how GMR affects model dynamics. Here, we test the effect of GMR by simulating the model then applying GMR on the dynamics of the model simulation. We do this by removing, via linear regression, the average time series taken across all nodes from each individual node. We show the effects of GMR on BEI, BTF, and Hopf model dynamics in Figure 7 (we exclude the noisy degree model because by construction only Gaussian noise will remain post GMR). For *G <* 2, coherent structures are removed, lowering VE1 and the majority of structure in the FC matrices. This dramatic removal of most WSDs persists for all *G* in the Hopf bifurcation model as the oscillations are roughly in phase and of similar frequency. This means that the Hopf model considered in Fig. 7 G,H is dominated by globally coherent signals at all coupling strengths of *G*.

**Fig. 7.**
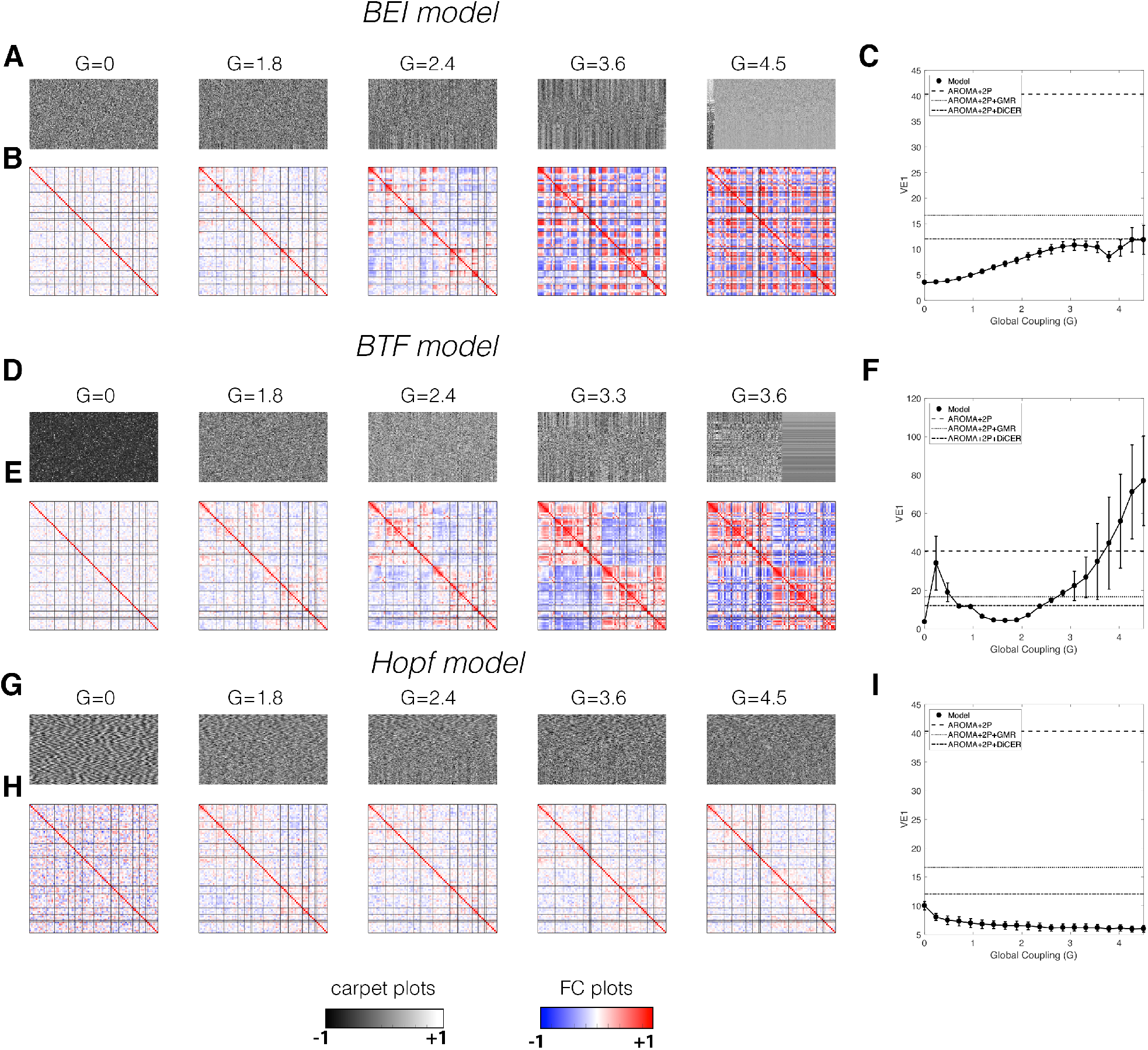
Model properties post GMR. Spatiotemporal carpet plots and FC matrices as a function of the global coupling parameter *G* in (A,B), (D,E) and (G,H) for one instance of the BEI, BTF, and Hopf models, respectively, after application of GMR. Variance explained by the first PC (VE1) is shown in (C), (F), and (I) for the BEI, BTF, and Hopf models post-GMR, respectively. The models were run 108 times and for 304s, mirroring the data. The curves for the model represent the mean metric values, with the error bars describing standard deviation at each *G*. The horizontal lines on the metrics (C,F,I) represent the mean VE1 under the three different denoising pipelines.

In the BTF model, we see the emergence of a “global” signal that appears in all nodes with a different delay - which is consistent with a travelling wave (52) - that will not be captured by the global mean used in GMR. Hence, for certain values of *G*, the BTF model post-GMR contains fewer WSDs than the BEI model (due to anticorrelated regions). The corresponding FC matrices also show some structure; forthe BTF model at *G* = 3.3, we see strong positive within-hemisphere FC and strongly anticorrelated inter-hemisphere FC. For the BEI model, positive and negative correlations are distributed more uniformly across edges. Neither of these resemble the network structure of empirical data post-GMR (Fig 4B).

VE1 for both the BEI (Fig. 7C) and Hopf models (Fig. 7I) is lower than the empirical VE1 for all values of *G*, indicating that this property of the data is not captured well by these models after the models have been subjected to GMR. The BTF model (Fig. 7F) shows a small parameter regime in which VE1 matches the empirical data (*G ≈*1.8 and *G ≈*2.4).

To summarise, we have shown that WSDs are present in model simulations as well as empirical rsfMRI data. The models do not contain structured non-neural signal (usually called noise in the rsfMRI context), suggests that WSDs are a real and an emergent property of neuronal dynamics unfolding on a connectome. However, globally synchronous WSDs can also be generated by relatively simple mechanisms, as captured in in the noisy degree model. This result suggests that the performance of dynamical models in fitting empirical data is primarily driven by fitting of first-order and relatively trivial globally synchronous dynamics rather than more complex, second-order properties that remain post-GMR.

### WSDs affect model fits to rsfMRI data

Having inspected the way in which WSDs affect resting-state fMRI data and models, we now turn to a more quantitative characterization of how they influence model fits to empirical data. We first focus on fitting the static FC architecture of the empirical resting-state fMRI data. In Figure 8A we show the results of fitting FC similarity, *R*_*s*_, in the BEI, BTF, Hopf and noisy degree models to the empirical data processed using three different denoising procedures. We find that model fits are higher following less aggressive preprocessing; i.e., for all except the BTF model, *R*_*s*_ is highest for ICA-AROMA+2P, followed by ICA-AROMA+2P+GMR, and lowest for ICA-AROMA+2P+DICER. The *R*_*s*_ values for data processed with ICA-AROMA+2P are generally within the state-of-the-art for rsfMRI modelling – exceeding 0.5. They fall below 0.4 when either GMR or DICER have been applied to the data and are lowest when GMR has been applied to both model and data. In the latter case, the maximal performance of all three DNMs drops: it reduces from 0.55 to 0.35 for the BEI model (30% drop), from 0.6 to 0.35 for the Hopf model (40% drop), and 0.56 to 0.1 in the Noisy degree model (82% drop). In these models, WSDs dominate the dynamics and their removal leaves residual dynamics that do not align with the empirical rsfMRI data. In other words, these models have a limited capacity to capture subtle dynamics that exist beyond widespread synchronization events.

**Fig. 8.**
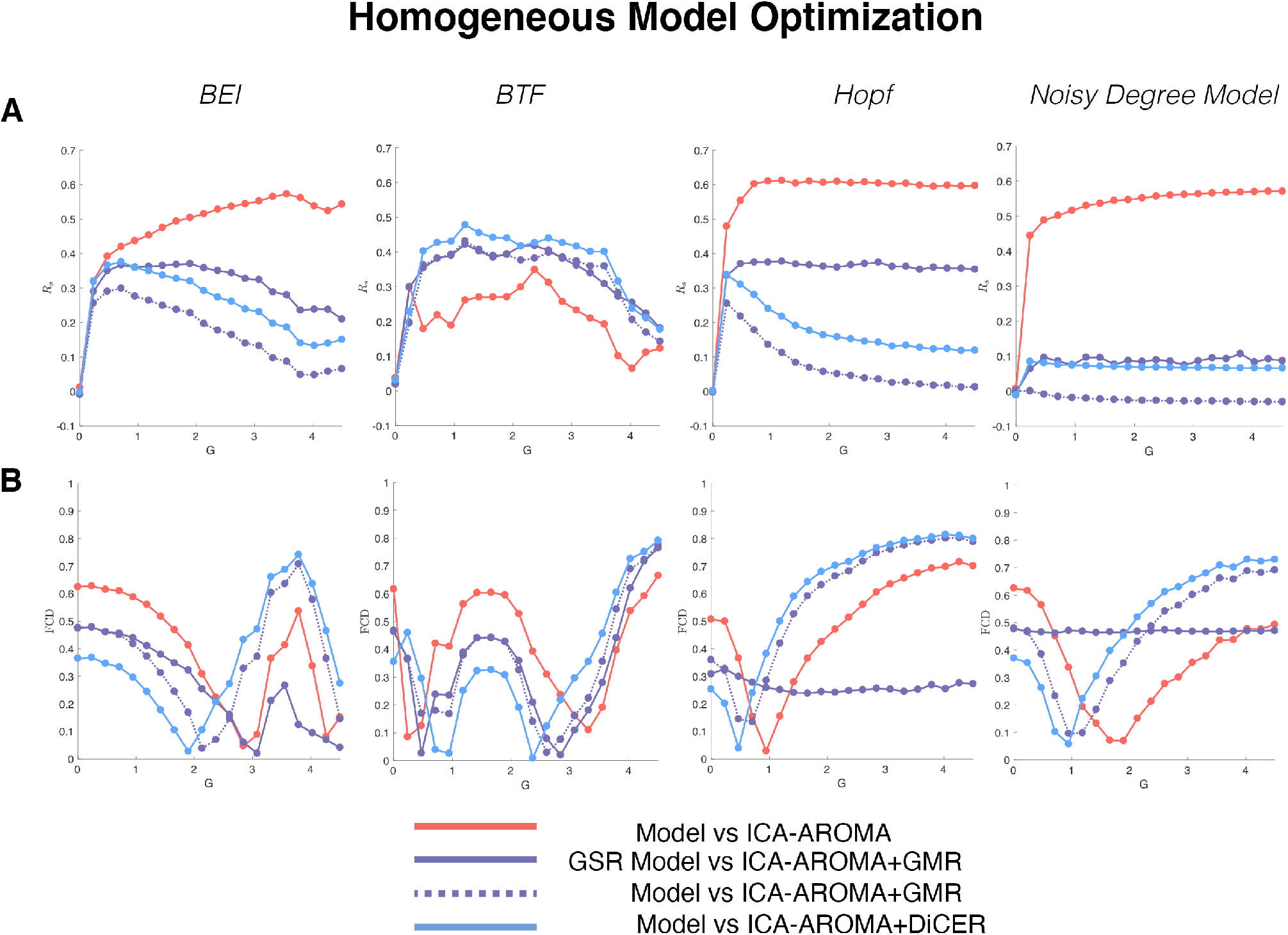
Data preprocessing influences fits of homogeneous models. (**A**) Similarity between model and empirical static FC, as quantified by *R*_*s*_. The simulated FC matrix is the mean over 108 runs of 302 seconds of simulated data. (**B**) Similarity of the spatiotemporal dynamics captured by the FCD statistic across all models. In both panels the comparisons are between the Model vs the ICA-AROMA+2P, GMR of the Model vs ICA-AROMA+2P+GMR, the Model vs ICA-AROMA+GMR, and the Model vs ICA-AROMA+DiCER in red, purple, dashed-purple and blue lines as indicated in the legend.

When fitting the data processed without correcting for global signals (i.e., ICA-AROMA+2P), the BEI model shows maximum *R*_*s*_ at *G >* 3 corresponding to regimes dominated by WSDs. In contrast, the Hopf and noisy degree models approach maximum *R*_*s*_ at *G <* 1. Maximal *R*_*s*_ are generally observed for *G <* 1.5 across all models when fitted to data processed using either GMR or DICER. Thus, as predicted, our analysis indicates that (1) model fits are mostly elevated in data with prominent WSDs compared to data with relatively fewer WSDs; and (2) the optimal working points of the models are obtained at higher values of *G* when fitting data with prominent WSDs. This analysis confirms that WSDs inflate model fit statistics. Critically, the noisy degree model, when fitted to data characterised by prominent WSDs (i.e., data that has not been subjected to GMR or DiCER), shows a comparable fit to the BEI and Hopf models, and a better fit than the BTF model. The noisy degree model only contains two parameters (*G, α*) as opposed to 32 in BTF and 21 in the BEI and 2 + *N* = 2 + 82 = 84 in Hopf (considering the frequencies are specified for each node). This result suggests that the FC matrix of the ICA-AROMA+2P-processed data can be explained simply by local noise modulated by regional variations in node degree and a common, global noise term.

Figure 5 also indicates that the dynamics in the BTF are weakly synchronised on average for *G <* 3.25 and thus, upon GMR, the fit of the empirical FC structure is slightly improved (Fig. 8, *R*_*s*_ *≈* 0.4). We note that while Fig. 5 shows a synchronised time series for *G >* 3.5, not all simulations reached this behaviour due largely to the chaotic behaviour of the BTF model, and it is possible that over longer time periods the system would synchronise.

One argument against the use of GMR in DNMs is that globally coherent signals are part of the dynamics, thus we then provide a comparison with data that has been de-noised with DiCER using models without GMR (Fig. 8A in blue). We see that in this case *R*_*s*_ is generally lower than the previous comparisons for the BEI, Hopf and the noisy degree model across all coupling strengths. The only exception again is the BTF where ICA-AROMA+2P+DiCER performs the best. Here again, since the BTF for *G <* 3.25 exhibits weakly coupled dynamics it is the regime most similar to the data processed under the ICA-AROMA+2P+DiCER stream (see examples in Fig. 2 and Fig. 3).

Figure 8B shows model fits to FCD. Unlike the *R*_*s*_ fits, there exists a minimum for FCD for nearly all models, with similar performance across models and datasets. The two exceptions to this trend are when GMR is applied to the Noisy degree model, where performance is expected by construction, and the Hopf model, as most of the dynamics in this model are in phase and thus substantially removed with GMR.

Critically, the noisy degree model exhibits similar FCD performance to the other three other models. This result is surprising, as it suggests that even the apparently complex spatiotemporal structure of time-resolved FC can be accounted for by local noise modulated by node degree and a common signal.

Given that FCD arrives at a global minimum for each model, minimising this metric can be used to choose the optimal value *G*. The corresponding optimised FC matrices are shown in Figure 9. The FC matrices summarise three key findings: (1) that for each model, the level of global correlation in the simulated FC matrices decreases in line with the degree to which pre-processing has removed WSDs from the fitted data (note that the scales change in the rows of Fig. 9); (2) The global coupling parameter *G* changes for each denoising pipeline and thus changes the level of coherence; and (3) GMR in any of the models changes FC dramatically, causing inter-hemispheric anti-correlations in the BTF and Hopf models, and large groups of anti-correlated networks in the BEI model. Together, these results indicate that both static and dynamic aspects of model-derived FC are sensitive to the degree to which WSDs are present in the fitted data.

**Fig. 9.**
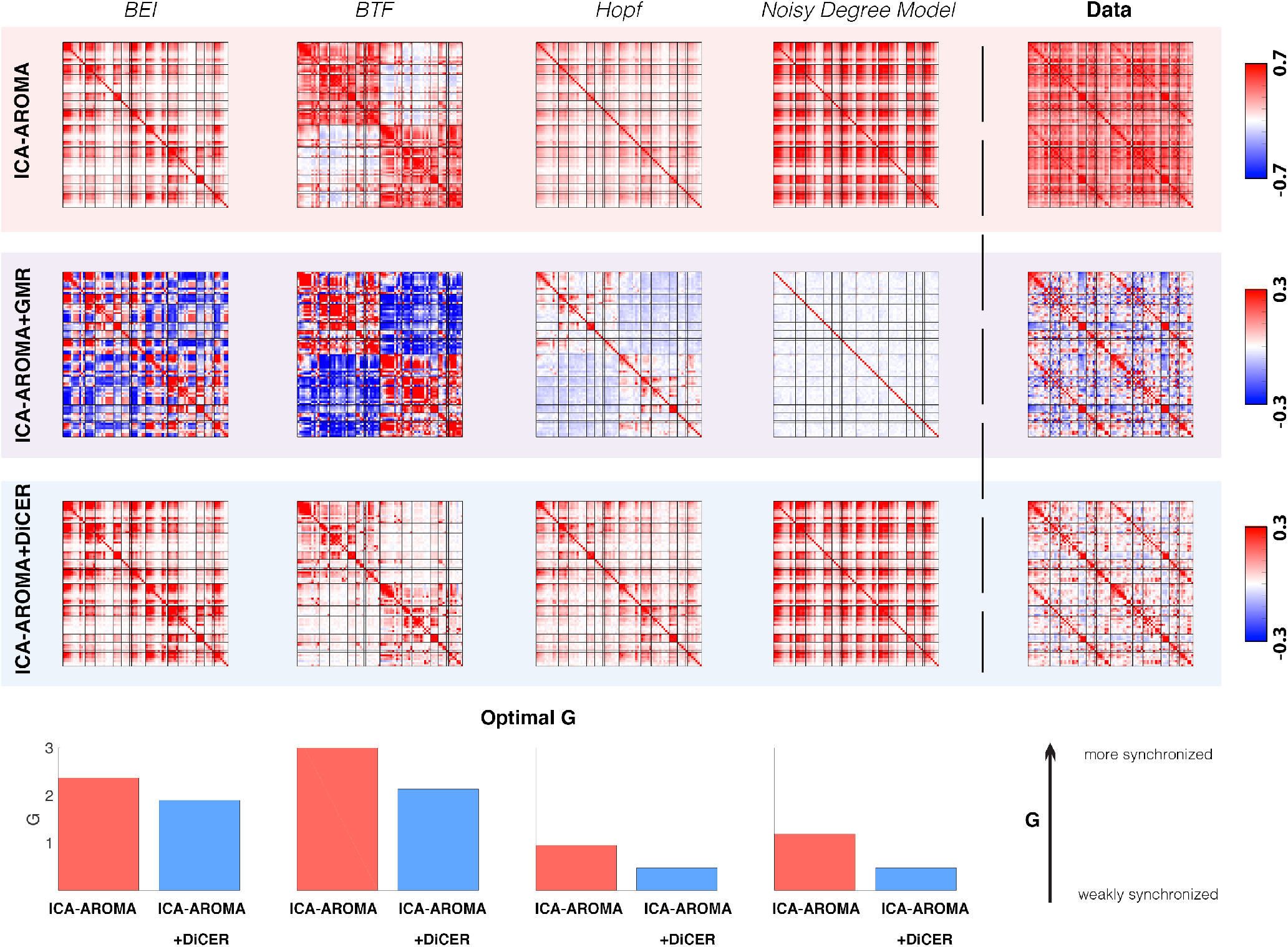
Model-derived FC matrices obtained after fitting data denoised at different levels. The FC matrices were generated by using the global coupling parameter, *G*, that minimised FCD across the three different denoising pipelines - ICA-AROMA+2P, ICA-AROMA+2P+GMR and ICA-AROMA+2P+DiCER. In the top and bottom rows the FC matrices were generated using Pearson correlations between pairs of regional time series without GMR and the middle row are the FC matrices post GMR. Here each FC matrix was a mean across 108 runs of data simulated 302 seconds, and the data was inspected so that no transients were included. Finally, the panels on the right replicate the FC matrices shown in Fig 4 as a point of comparison, i.e. the average over 108 subjects. The histograms at the bottom reflect the *G* estimated from model optimisation for ICA-AROMA+2P, and ICA-AROMA+2P+DiCER. Note ICA-AROMA+2P+GMR was omitted as both the Hopf and the Noisy degree model have no clear FCD minima.

### Heterogeneous network models

The results presented thus far indicate that the DNMs considered here show weak performance (with respect to *R*_*s*_ for denoised data) with regards to fitting the static or dynamic FC properties of thoroughly denoised rsfMRI data (i.e., data processed with GMR or DiCER). Although the models perform well in data that have not undergone extensive removal of WSDs (i.e., data processed with ICA-AROMA+2P), the noisy degree model performs similarly well, suggesting that the multi-parameter, non-linear, and biophysically-informed DNMs dynamical models themselves are mostly capturing very basic global synchronization properties of fMRI dynamics. Here, we explore whether adding additional degrees of freedom to a DNM can improve model performance (34, 36, 37). We incorporate heterogeneity either at the node level by adjusting local population dynamics, or at the edge level by modifying node-to-node connection weights, as described in the Methods. We focus on heterogeneity in the Hopf model for computational efficiency.

#### Node heterogeneity

In the first instance, we use the Hopf model and fit bifurcation parameters *a*_*i*_ to match the local power spectrum of the model to the data (see Methods for model optimization). The results of the optimization are shown in Figure 10 and indicate that, as with the homogeneous model, the node-heterogeneous Hopf model shows *R*_*s*_ = 0.68 for ICA-AROMA+2P data (achieved at a lower value of *G* than the homogeneous model; Fig. 10 A), but much poorer fits to the data processed with GMR (Fig. 10 B) or DiCER (Fig. 10 C). In contrast, FCD is fitted slightly better in the data processed with GMR and DICER. Regardless of how the data were processed, we find that the fitted bifurcation *a*_*i*_ parameters have a relatively similar distribution across nodes at the optimal *G* (by choosing the minimal FCD Fig. 10 D). Only some some nodes change from noisy to oscillatory (or vice versa) in the insula, frontal, cingulate cortex. Such variations can lead to dramatically different conclusions about regional contributions to spontaneous BOLD dynamics. Taken together, these results indicate that the introduction of node heterogeneity to the Hopf model yields no appreciable benefit (*R*_*s*_ = 0.68, 0.4, 0.28 vs *R*_*s*_ = 0.61, 0.38, 0.32 in the homogeneous variant of the Hopf model when fitted to the ICA-AROMA+2P,ICA-AROMA+2P+GMR, and ICA-AROMA+2P+DiCER pipelines respectively), particularly in aggressively denoised data. This lack of improvement in performance arises despite the additional complexity of 88 new free parameters optimized within-sample.

**Fig. 10.**
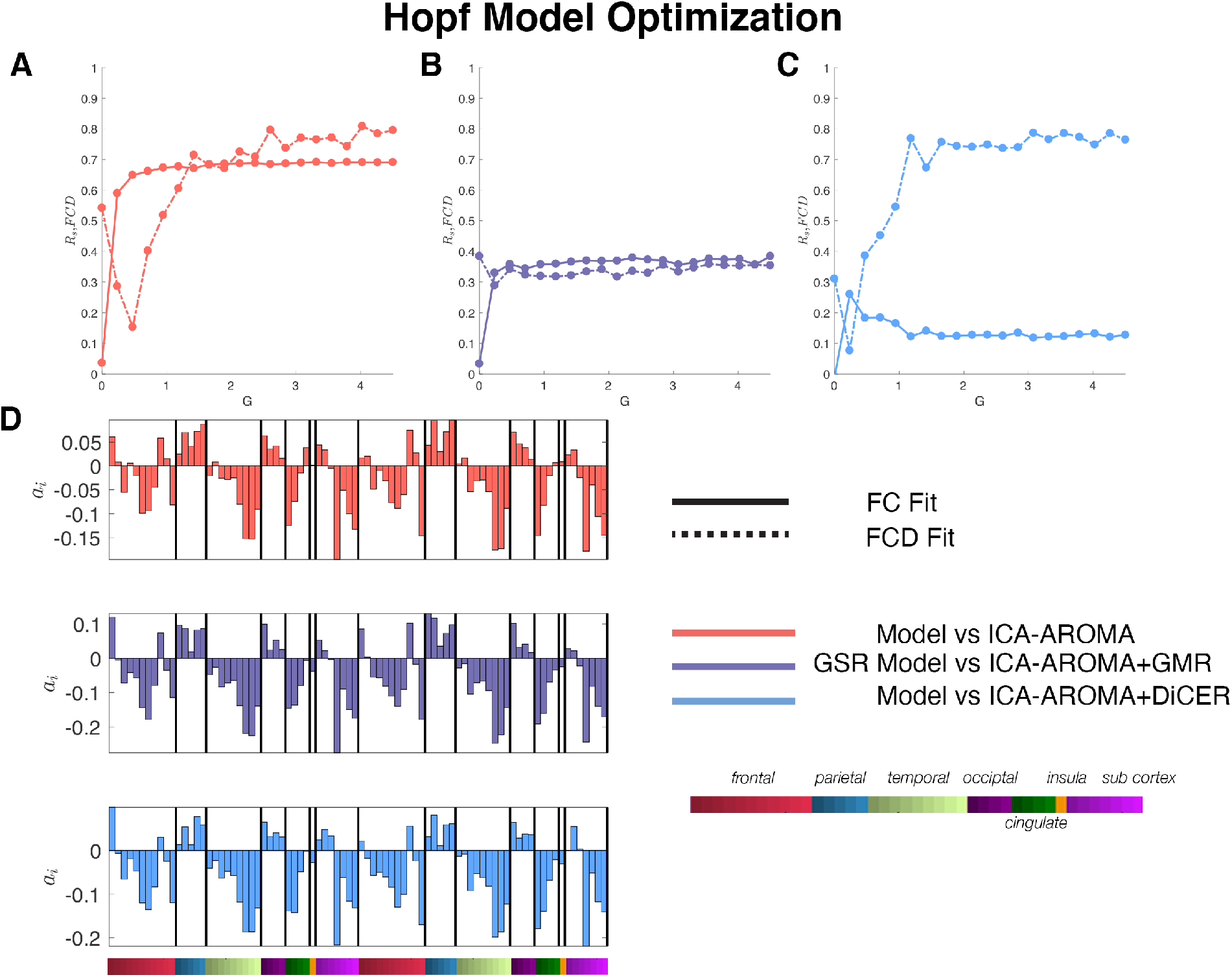
Performance of the node-heterogeneous Hopf model. **A**,**B**,**C** show model fits with *R*_*s*_ or FCD for the Hopf model optimized to match the local power spectrum of ICA-AROMA+2P, ICA-AROMA+2P+GMR or ICA-AROMA+2P+DiCER respectively. The dashed lines indicate FCD and the solid lines indicate *R*_*s*_. Fit statistics were generated by fitting the data and over 108 runs of simulations of 302 seconds to match the data that is being fit. **D** indicates the local bifurcation parameters *a*_*i*_ found after optimization to the three pipelines which are coloured according to the legend on the right. The local bifurcation parameters are grouped according to major regions specified indicated by the colours below and named in the legend on the right.

#### Edge heterogeneity

Finally, we introduce heterogeneity to the Hopf model at the edge level by modifying the connections between nodes in order to estimate the effective connectivity *E*_*ij*_ which is a method that has been used in DCM (92) and in other contexts rsfMRI (69). This optimization results in using *E*_*ij*_ (which is a modified *C*_*ij*_) in place of the structural connectivity to simulate the homogeneous Hopf bifurcation model (i.e. with *a*_*i*_ uniform across all nodes). As described in the methods, this procedure modifies existing connections in *C*_*ij*_ to force the dynamic coupling between two regions in the simulations to match the existing data. As also indicated in the methods, since a large number of parameters are being optimized – i.e. all the edges, which in out case is 3828 parameters – this procedure is prone to over-fitting. One solution to mitigate over-fitting is to estimate model parameters in a training set and evaluate model fits in an independent test set. Here, we estimate *E*_*ij*_ by optimizing the fits to *FCD* on 80% (*n* = 86) of the data and test it on the remaining 20% (*n* = 22) of individuals. Once estimated, the model is run 22 times for 302 seconds to match the testing data. This cross validation is repeated 108 times where the data is split into test and retest randomly without replacement. We focus on fitting data that has undergone GSR or DiCER, since these are the datasets that are not adequately modelled by the homogeneous or node-heterogeneous models.

The results of our cross-validated optimization procedure are shown in Figure 11. For data processed with ICA-AROMA+2P+GMR (Fig. 11A), *R*_*s*_ doubles from 0.4 in the homogeneous case to 0.8 in the edge-heterogeneous case. However, FCD shows a relatively poor fit, with the best value reaching only 0.3, which is one of the worst performing amongst nearly all model variants. This is again likely due to the fact that within the Hopf model, the coupled oscillations are in phase even though the optimized *E*_*ij*_ increases the weights of most of the connections (Fig. 11C). We note that although *R*_*s*_ is high, the overall level of FC across pairs of nodes tends to be lower in the model compared to data (Fig. 11B choosing the max *R*_*s*_ for *G*) with data.

**Fig. 11.**
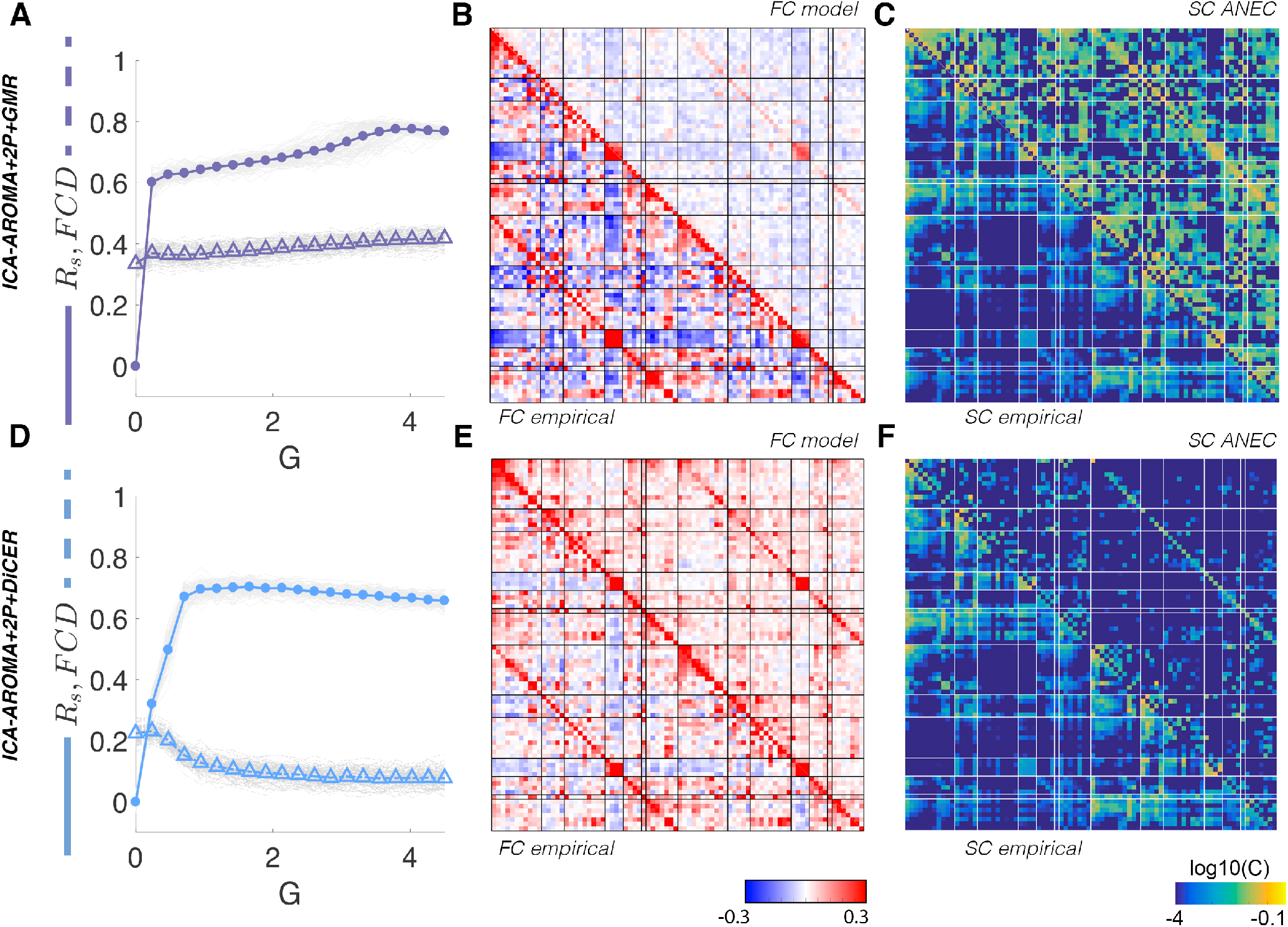
Performance of the edge-heterogeneous Hopf model. **A**,**D** Optimization plots showing *R*_*s*_ with solid lines and FCD with dashed lines for simulations optimized to fit data processed with ICA-AROMA+2P+GMR and ICA-AROMA+2P+DiCER, respectively. Each grey solid and dashed line represents the calculation of *R*_*s*_ and FCD, respectively, from a single estimate derived from the cross validation procedure. The coloured lines represent the mean optimization plots over the 108 iterations of the cross validation procedured described in the methods. Panels **B** and **E** show optimal FC matrices with the corresponding empirical training sets for using the ICA-AROMA+2P+GMR and ICA-AROMA+2P+DiCER pipelines respectively

For data processed using ICA-AROMA+2P+DiCER, optimizing *E*_*ij*_ doubles *R*_*s*_ from *≈* 0.35 (in the homogeneous case) to *≈* 0.75. We also find that FCD is fitted reasonably well across a broad range of *G*. Visually comparing the FC matrix at the chosen minimum of *G* = 2.5 we observe that the structure (Fig. 11E) of the estimated model is similar to the data, and spans a similar range of FC values. On comparisons to the carpet plots and VE1 in Figure 12 we also find that the model visually replicates the fluctuations present in the data.

**Fig. 12.**
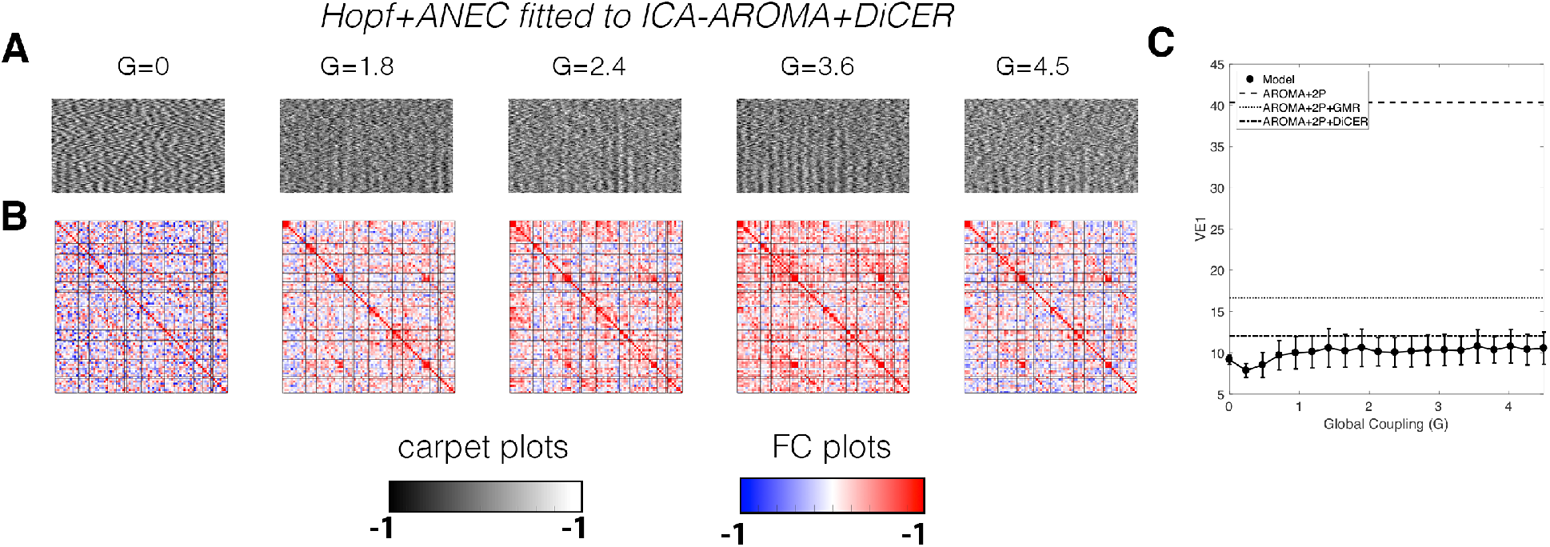
WSDs in the resulting Hopf+ANEC model optimised for the ICA-AROMA+2P+DiCER pipeline. (A) carpet plots and (B) functional connectivity matrices as a function of the global coupling parameter *G*. (C) Variance explained by the first PC (VE1), with solid line indicating the mean VE1 across one iteration of the test-retest cross validation fold, i.e., 22 runs simulated at 304s with the error bars representing the standard deviation of these runs. Horizontal lines represent the mean VE1 under the three different denoising pipelines. In panel C, the re-test sample was on one iteration trained from 80% of the data.

These results suggest that introducing heterogeneity at the edge level offers a promising approach to fitting data with WSDs removed, and that this can be done within a cross-validated framework that minimizes the potential for over-fitting.

## Discussion

Whole-brain modeling of rsfMRI data with DNMs has been used to understand the basic principles of spontaneous neural dynamics (23, 25). However, ongoing debate over the degree to which such data are contaminated by noise, and in particular the origins of prominent WSDs, raises questions about the specific data features that are accurately captured by DNMs (42, 88). Here, we investigated the relationship between data quality and model performance and found that the level of WSDs within a dataset influences the fits and dynamic regime of the models (Fig. 8; Fig. 5). For data with sufficiently large WSDs, a simple two parameter linear model performs just as well as complex, non-linear and multi-parameter DNMs (Fig., 6,8. Finally, we present evidence that model heterogeneity, particularly at the level of edges, can improve model fits regardless of how the data were processed, at the expense of increased model complexity (Fig. 11).

### How WSDs bias models

Quality control of rsfMRI data has become more sophisticated in recent years, involving extensive reporting of various metrics and visualisations attainable at the level of individual subjects (44, 54, 87) our group summaries (41, 43). These measures have been used extensively to inform the development of preprocessing pipelines for experimental analyses of rsfMRI data (40, 41), but are seldom considered in the modelling literature.

Our analysis focused specifically on how WSDs affect model performance. The degree to which WSDs in empirical rsfMRI represent neuronal or non-neuronal processes is an important, unresolved question. Growing evidence indicates that many WSDs are tied to head motion and respiratory variations (42, 44). There is also evidence that at least some WSDs may be neuronal in origin (48, 58), and such WSDs would be compatible with widespread neuromodulatory inputs (93). DNMs contain no noise. Thus, the emergence of WSDs in these models, for sufficiently high *G*, suggests that WSDs can indeed arise as an intrinsic property of the dynamics unfolding on the underlying SC matrix. However, the emergence of globally synchronised dynamics in the model depends on the level of *G*, and this level is chosen to fit empirical data. Our results indicate that the optimal level of *G* for a given model depends strongly on how the data are preprocessed. Fitting models to empirical properties such as *R*_*s*_ and *FCD* estimated from data with prominent WSDs (e.g., without prior application of GMR or DiCER) pushes the models to a highly synchronised, high-*G* regime, in which prominent WSDs emerge in the model time series. For data with WSDs removed (i.e., following GMR and DiCER), the models are pushed to a weakly synchronised, low-*G* regime. Applying GMR to both the model and data results in a poor fit, as the residual synthetic and empirical FC properties differ markedly. These results suggest that while DNMs, particularly the Hopf and BEI models accurately fit “first order” properties of globally coherent signals, they do a poor job of capturing “second order”, more complex aspects of spatiotemporal structure in the data. Since, the noisy degree model fits these first order properties just as well as the complex, multi-parameter DNMs, such properties are essentially trivial from a modelling perspective. Our findings thus draw attention to the need for caution when interpreting model fits to empirical rsfMRI data. Critically, these results are not just a consequence of GMR altering the distribution of FC values (89, 94) since we obtain similar findings with DiCER-processed data.

Some of these issues likely stem from the uniformity of node parameters, as the nodes in the absence of coupling will have the same natural frequency. As such, high values of *G* will force all to align in phase, resulting in globally coherent activity. One way to circumvent this behaviour while retaining uniform node parameters is to introduce time delays to disrupt in-phase coherence. Another approach is to use nonlinear chaotic oscillations such as those in the BTF. Notably, this model performed better in data subjected to more aggressive denoising (Fig 8).

A third way to combat the dominating effect of WSDs in model fitting is to introduce heterogeneity at the level of either nodes or edges to force node dynamics with different natural frequencies (36) and/or to modify connections to reduce the prevalence of globally coherent fluctuations. We presented preliminary evidence to suggest that edge-heterogeneity may represent a promising avenue forward, as our edge-heterogeneous Hopf model dramatically improved both *R*_*s*_ and *FCD* when compared to the homogeneous model fitted to data processed with the aggressive ICA-AROMA+2P+DiCER pipeline (Fig. 11). The relatively poor performance of our node-heterogeneous Hopf model (Fig. 10) may be due our reliance on a specific type of optimisation, or because the heterogeneity of the bifurcation parameter is not sufficient to break the global synchrony present in the Hopf model (c.f. Fig 5 with Fig 7). Studying the effect of alternative, physiologically-grounded methods for introducing heterogeneity ((36, 38)), will be an important extension of this work.

### The noisy degree model

In this study we showed, by analysing carpet plots and model equations, that a simple model of global signal modulation is enough to produce the properties of *R*_*s*_ and *FCD* when comparing data processed with the ICA-AROMA+2P pipeline. This is a surprising result, and calls into question the utility of highly complex, multiparameter biophysical models. We propose that a fruitful way forward in model development and evaluation will involve first testing the performance of any new model against simple benchmarks such as the noisy degree model or other simple linear approximations (7).

One important implication of the strong performance of the noisy degree model in data processed with the ICA-AROMA+2P pipeline is that signal dynamics in such data (i.e., data in which no explicit correction for WSDs has been performed) are predominantly explained by a shared common signal, local noise, and node degree of the SC matrix. The model simply captures the fact that a given region’s activity will be more strongly driven by the common signal if it is highly connected to other areas.

### Limitations

Our analyses focused on a single, typical quality fMRI dataset and we only considered a single brain parcellation. Empirical properties of the data (95, 96) and model performance (35) can depend on parcel resolution, but our primary results should generalise (61) as they describe intrinsic properties of the DNMs themselves and the rsfMRI data features we consider (e.g., complex WSD structure) are observed in diverse datasets and across different parcellation resolutions (e.g., (43)).

We evaluated relative model performance qualitatively. A more formal treatment would include measures of model fit that account for model complexity, such as those used in dynamic causal modelling (97). However, these are not routinely applied in the large-scale DNM literature.

Finally, we simulated model dynamics on an SC matrix generated from a cohort of individuals that differed to the fMRI dataset. While this is common practice in the DNM literature, model fitting can be done at the single subject level given concurrent SC and FC measures ((7)). Future work may also consider the effects of diffusion MRI preprocessing variations (98) on model results.

## Conclusions

We show that the results of modelling studies depend strongly on the way in which the experimental data have been preprocessed. We hope that our findings will encourage greater communication between theorists and experimenters, to ensure that any proposed models are capturing important properties of the data, rather than low-order (neuronal or non-neuronal) structure. A better understanding of large-scale brain dynamics will be contingent on a tight integration between theory and experiment.

## Appendix

### Parameters for the models

This section contains the numerical parameters for the dynamic models.

### Numerical approximation for *J*_*i*_

In the BEI model, a key step is determining the approximation for the inhibitory parameter *J*_*i*_ that ensures that the firing rate at each node *i* has excitatory firing rate of 3 Hz. As described in Deco et al. (26), the value of *J*_*i*_ is numerically solved via a Greedy search algorithm, where the model is estimated for 10 seconds, then at each node the value of *J*_*i*_ is adjusted by a factor Δ*J*, which is dependent on the mean firing rate at node *i*. If the firing rate is above 3 Hz, *J*_*i*_ = *J*_*i*_ + Δ, if below 3 Hz, *J*_*i*_ = *J*_*i*_ *−* Δ*J*, where Δ*J* is additionally reduced until the node is within the firing rate from 2.63*− −*3.55 Hz. To improve convergence times, an analytic form is first used as an initial starting point by finding the fixed point of the dynamical system i.e. the condition that 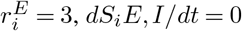 and setting *ν*_*i*_(*t*) = 0.

This yields the analytical approximation for *J*_*i*_ as:

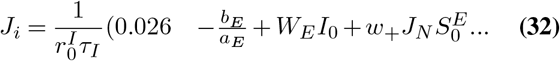

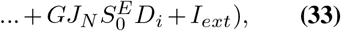

where

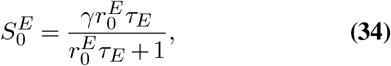

and 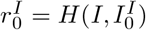, where 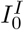 is calculated by solving the following transcendental equation:

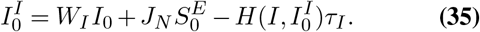

We note that the balanced inhibitory criterion is related to node degree *D*_*i*_ by the following relationship:

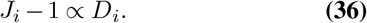

### BOLD forward model - The Balloon model

Here we include the hemodynamic equations for the balloon model (28, 79) used as a forward BOLD model in this study. For a neuronal input at node *i, z*^*i*^(*t*), the following dynamical equations detail the resulting fractional changes in blood flow *f*_*i*_(*t*), blood volume *v*_*i*_(*t*) and deoxygenated hemoglobin content *q*_*i*_(*t*):

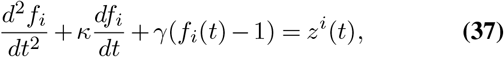

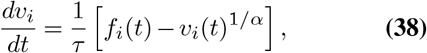

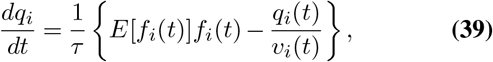

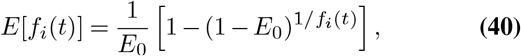

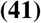

where *E* is the oxygen extraction fraction, *E*_0_ is the resting oxygenation fraction *τ* is the mean hemodynamic transit time, *α* is the Grubb exponent, *γ* is the flow elimination constant, and *κ* is the flow decay rate. The hemodynamic changes in *v*_*i*_ and *q*_*i*_ drive the BOLD response *y*_*i*_(*t*) which is modelled by the semi-empirical relationship (99):

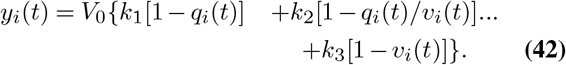

where the parameters *k*_1_, *k*_2_, *k*_3_ depend on the field strength, and acquisition. The parameter values are detailed in the table below:

## ACKNOWLEDGEMENTS

AF was supported by the Australian Research Council (ID: FT130100589), National Health and Medical Research Council (NHMRC; ID: 1104580), and the Sylvia and Charles Viertel Charitable Foundation. This work was supported by the MASSIVE HPC facility (www.massive.org.au). The authors thank Stuart Heitmann and James Roberts for support of the Brain dynamics toolbox, and Christopher honey and Olaf Sporns for sharing code.

## Bibliography

1. B. Biswal, F. Zerrin Yetkin, V. M. Haughton, and J. S. Hyde. Functional connectivity in the motor cortex of resting human brain using echo-planar mri. Magnetic resonance in medicine 34, 537 (1995).

2. A. Fornito and E. T. Bullmore. What can spontaneous fluctuations of the blood oxygenation-level-dependent signal tell us about psychiatric disorders? Current opinion in psychiatry 23, 239 (2010).

3. M. D. Fox and M. E. Raichle. Spontaneous fluctuations in brain activity observed with functional magnetic resonance imaging. Nature reviews neuroscience 8, 700 (2007).

4. M. D. Fox, A. Z. Snyder, J. L. Vincent, et al. The human brain is intrinslly organized into dynamic, anticorrelated functional networks. Proc. Natl. Acad. Sci. USA 102, 9673 (2005).

5. S. M. Smith, P. T. Fox, K. L. Miller, et al. Correspondence of the brain’s functional architecture during activation and rest. Proceedings of the National Academy of Sciences 106, 13040 (2009).

6. C. F. Beckmann and S. M. Smith. Probabilistic independent component analysis for functional magnetic resonance imaging. IEEE transactions on medical imaging 23, 137 (2004).

7. A. Messé, D. Rudrauf, H. Benali, and G. Marrelec. Relating structure and function in the human brain: relative contributions of anatomy, stationary dynamics, and non-stationarities. PLoS computational biology 10 (2014).

8. P. Hagmann, L. Cammoun, X. Gigandet, et al. Mapping the structural core of human cerebral cortex. PLoS biology 6 (2008).

9. J. L. Vincent, G. H. Patel, M. D. Fox, et al. Intrinsic functional architecture in the anaesthetized monkey brain. Nature 447, 83 (2007).

10. V. Zerbi, M. Wiesmann, T. L. Emmerzaal, et al. Resting-state functional connectivity changes in aging apoe4 and apoe-ko mice. Journal of Neuroscience 34, 13963 (2014).

11. V. Zerbi, J. Grandjean, M. Rudin, and N. Wenderoth. Mapping the mouse brain with rs-fmri: An optimized pipeline for functional network identification. Neuroimage 123, 11 (2015).

12. E. S. Finn, X. Shen, D. Scheinost, et al. Functional connectome fingerprinting: identifying individuals using patterns of brain connectivity. Nature neuroscience 18, 1664 (2015).

13. E. Amico and J. Goñi. The quest for identifiability in human functional connectomes. Scientific reports 8, 8254 (2018).

14. S. M. Smith, T. E. Nichols, D. Vidaurre, et al. A positive-negative mode of population covariation links brain connectivity, demographics and behavior. Nature neuroscience 18, 1565 (2015).

15. J. Li, R. Kong, R. Liegeois, et al. Global signal regression strengthens association between resting-state functional connectivity and behavior. BioRxiv p. 548644 (2019).

16. A. Fornito, A. Zalesky, D. S. Bassett, et al. Genetic influences on cost-efficient organization of human cortl functional networks. Journal of Neuroscience 31, 3261 (2011).

17. D. C. Glahn, A. Winkler, P. Kochunov, et al. Genetic control over the resting brain. Proceedings of the National Academy of Sciences 107, 1223 (2010).

18. M. D. Fox, A. Z. Snyder, J. M. Zacks, and M. E. Raichle. Coherent spontaneous activity accounts for trial-to-trial variability in human evoked brain responses. Nature neuroscience 9, 23 (2006).

19. M. D. Fox, A. Z. Snyder, J. L. Vincent, and M. E. Raichle. Intrinsic fluctuations within cortical systems account for intertrial variability in human behavior. Neuron 56, 171 (2007).

20. D. Zhang and M. E. Raichle. Disease and the brain’s dark energy. Nature Reviews Neurology 6, 15 (2010).

21. A. Abraham, M. P. Milham, A. Di Martino, et al. Deriving reproducible biomarkers from multi-site resting-state data: An autism-based example. NeuroImage 147, 736 (2017).

22. J. A. Hadley, N. V. Kraguljac, D. M. White, et al. Change in brain network topology as a function of treatment response in schizophrenia: a longitudinal resting-state fmri study using graph theory. npj Schizophrenia 2, 1 (2016).

23. G. Deco, V. K. Jirsa, P. A. Robinson, M. Breakspear, and K. Friston. The dynamic brain: from spiking neurons to neural masses and cortical fields. PLoS computational biology 4 (2008).

24. J. Alstott, M. Breakspear, P. Hagmann, L. Cammoun, and O. Sporns. Modeling the impact of lesions in the human brain. PLoS computational biology 5 (2009).

25. M. Breakspear. Dynamic models of large-scale brain activity. Nature neuroscience 20, 340 (2017).

26. G. Deco, A. Ponce-Alvarez, P. Hagmann, et al. How local excitation–inhibition ratio impacts the whole brain dynamics. Journal of Neuroscience 34, 7886 (2014).

27. K. E. Stephan, N. Weiskopf, P. M. Drysdale, P. A. Robinson, and K. J. Friston. Comparing hemodynamic models with dcm. Neuroimage 38, 387 (2007).

28. R. B. Buxton, E. C. Wong, and L. R. Frank. Dynamics of blood flow and oxygenation changes during brain activation: the balloon model. Magnetic resonance in medicine 39, 855 (1998).

29. P. Sanz Leon, S. A. Knock, M. M. Woodman, et al. The virtual brain: a simulator of primate brain network dynamics. Frontiers in neuroinformatics 7, 10 (2013).

30. X.-J. Wang and H. Kennedy. Brain structure and dynamics across scales: in search of rules. Current opinion in neurobiology 37, 92 (2016).

31. R. Chaudhuri, K. Knoblauch, M.-A. Gariel, H. Kennedy, and X.-J. Wang. A large-scale circuit mechanism for hierarchical dynamical processing in the primate cortex. Neuron 88, 419 (2015).

32. P. Kale, A. Zalesky, and L. L. Gollo. Estimating the impact of structural directionality: How reliable are undirected connectomes? Network Neuroscience 2, 259 (2018).

33. M. Breakspear, J. R. Terry, and K. J. Friston. Modulation of excitatory synaptic coupling facilitates synchronization and complex dynamics in a biophysical model of neuronal dynamics. Network: Computation in Neural Systems 14, 703 (2003).

34. G. Deco, M. L. Kringelbach, V. K. Jirsa, and P. Ritter. The dynamics of resting fluctuations in the brain: metastability and its dynamical cortical core. Scientific reports 7, 1 (2017).

35. C. Honey, O. Sporns, L. Cammoun, et al. Predicting human resting-state functional connectivity from structural connectivity. Proceedings of the National Academy of Sciences 106, 2035 (2009).

36. M. Demirtaş, J. B. Burt, M. Helmer, et al. Hierarchical heterogeneity across human cortex shapes large-scale neural dynamics. Neuron 101, 1181 (2019).

37. P. Wang, R. Kong, X. Kong, et al. Inversion of a large-scale circuit model reveals a cortical hierarchy in the dynamic resting human brain. Science advances 5, eaat7854 (2019).

38. M. L. Deco Gustavo and, A. Arnatkeviciute, S. Oldham, et al. Dynamical consequences of regional heterogeneity in the brains transcriptional landscape. bioRxiv (2021).

39. P. Sanz-Leon, P. A. Robinson, S. A. Knock, et al. Nftsim: theory and simulation of multiscale neural field dynamics. PLoS computational biology 14, e1006387 (2018).

40. R. Ciric, D. H. Wolf, J. D. Power, et al. Benchmarking of participant-level confound regression strategies for the control of motion artifact in studies of functional connectivity. Neuroimage 154, 174 (2017).

41. L. Parkes, B. D. Fulcher, M. Yücel, and A. Fornito. An evaluation of the effcy, reliability, and sensitivity of motion correction strategies for resting-state functional MRI. NeuroImage 171, 415 (2018).

42. J. D. Power, M. Plitt, T. O. Laumann, and A. Martin. Sources and impltions of whole-brain fmri signals in humans. Neuroimage 146, 609 (2017).

43. K. M. Aquino, B. D. Fulcher, L. Parkes, K. Sabaroedin, and A. Fornito. Identifying and removing widespread signal deflections from fmri data: Rethinking the global signal regression problem. NeuroImage 212, 116614 (2020).

44. J. D. Power, M. Plitt, S. J. Gotts, et al. Ridding fmri data of motion-related influences: Removal of signals with distinct spatial and physical bases in multiecho data. Proceedings of the National Academy of Sciences 115, E2105 (2018).

45. G. H. Glover, T.-Q. Li, and D. Ress. Image-based method for retrospective correction of physiologl motion effects in fmri: Retroicor. Magnetic Resonance in Medicine: An Official Journal of the International Society for Magnetic Resonance in Medicine 44, 162 (2000).

46. P. Kundu, N. D. Brenowitz, V. Voon, et al. Integrated strategy for improving functional connectivity mapping using multiecho fmri. Proceedings of the National Academy of Sciences 110, 16187 (2013).

47. J. D. Power. Temporal has not properly separated global fmri signals: A comment on glasser et al.(2018). Neuroimage (2019).

48. M. F. Glasser, T. S. Coalson, J. D. Bijsterbosch, et al. Using temporal ica to selectively remove global noise while preserving global signal in functional mri data. NeuroImage 181, 692 (2018).

49. J. D. Power. A simple but useful way to assess fmri scan qualities. Neuroimage 154, 150 (2017).

50. J. Cabral, E. Hugues, O. Sporns, and G. Deco. Role of local network oscillations in restingstate functional connectivity. Neuroimage 57, 130 (2011).

51. C. J. Honey, R. Kötter, M. Breakspear, and O. Sporns. Network structure of cerebral cortex shapes functional connectivity on multiple time scales. Proceedings of the National Academy of Sciences 104, 10240 (2007).

52. J. A. Roberts, L. L. Gollo, R. G. Abeysuriya, et al. Metastable brain waves. Nature communications 10, 1 (2019).

53. R. A. Poldrack, E. Congdon, W. Triplett, et al. A phenome-wide examination of neural and cognitive function. Scientific data 3, 160110 (2016).

54. O. Esteban, C. J. Markiewicz, R. W. Blair, et al. fmriprep: a robust preprocessing pipeline for functional mri. Nature methods 16, 111 (2019).

55. V. Fonov, A. Evans, R. McKinstry, C. Almli, and D. Collins. Unbiased nonlinear average age-appropriate brain templates from birth to adulthood. NeuroImage 47, Supplement 1, S102 (2009).

56. R. H. R. Pruim, M. Mennes, D. van Rooij, et al. Ica-AROMA: A robust ICA-based strategy for removing motion artifacts from fmri data. NeuroImage 112, 267 (2015).

57. J. D. Power, A. Mitra, T. O. Laumann, et al. Methods to detect, characterize, and remove motion artifact in resting state fmri. NeuroImage 84, 320 (2014).

58. M. F. Glasser, T. S. Coalson, J. D. Bijsterbosch, et al. Classification of temporal ica components for separating global noise from fmri data: Reply to power. NeuroImage (2019).

59. R. S. Desikan, F. Ségonne, B. Fischl, et al. An automated labeling system for subdividing the human cerebral cortex on mri scans into gyral based regions of interest. Neuroimage 31, 968 (2006).

60. A. M. Dale, B. Fischl, and M. I. Sereno. Cortical surface-based analysis: I. segmentation and surface reconstruction. NeuroImage 9, 179 (1999).

61. D. C. Van Essen, S. M. Smith, D. M. Barch, et al. The wu-minn human connectome project: an overview. Neuroimage 80, 62 (2013).

62. M. F. Glasser, S. N. Sotiropoulos, J. A. Wilson, et al. The minimal preprocessing pipelines for the human connectome project. NeuroImage 80, 105 (2013).

63. J.-D. Tournier, R. Smith, D. Raffelt, et al. Mrtrix3: A fast, flexible and open software framework for medical image processing and visualisation. NeuroImage 202, 116137 (2019).

64. J. D. Tournier, F. Calamante, and A. Connelly. Improved probabilistic streamlines tractography by 2nd order integration over fibre orientation distributions. In Proceedings of the international society for magnetic resonance in medicine, volume 1670. Ismrm (2010).

65. J.-D. Tournier, F. Calamante, and A. Connelly. Robust determination of the fibre orientation distribution in diffusion mri: non-negativity constrained super-resolved spherical deconvolution. Neuroimage 35, 1459 (2007).

66. J.-D. Tournier, F. Calamante, and A. Connelly. Mrtrix: diffusion tractography in crossing fiber regions. International journal of imaging systems and technology 22, 53 (2012).

67. R. E. Smith, J.-D. Tournier, F. Calamante, and A. Connelly. Anatomically-constrained tractography: improved diffusion mri streamlines tractography through effective use of anatomical information. Neuroimage 62, 1924 (2012).

68. R. E. Smith, J.-D. Tournier, F. Calamante, and A. Connelly. The effects of sift on the reproducibility and biological accuracy of the structural connectome. Neuroimage 104, 253 (2015).

69. M. Gilson, R. Moreno-Bote, A. Ponce-Alvarez, P. Ritter, and G. Deco. Estimation of directed effective connectivity from fmri functional connectivity hints at asymmetries of cortical connectome. PLoS computational biology 12 (2016).

70. K.-F. Wong and X.-J. Wang. A recurrent network mechanism of time integration in perceptual decisions. Journal of Neuroscience 26, 1314 (2006).

71. C. Morris and H. Lecar. Voltage oscillations in the barnacle giant muscle fiber. Biophysical journal 35, 193 (1981).

72. L. L. Gollo, C. Mirasso, O. Sporns, and M. Breakspear. Mechanisms of zero-lag synchronization in cortical motifs. PLoS Comput Biol 10, e1003548 (2014).

73. S. Heitmann, M. J. Aburn, and M. Breakspear. The brain dynamics toolbox for matlab. Neurocomputing 315, 82 (2018).

74. F. Freyer, J. A. Roberts, R. Becker, et al. Biophysical mechanisms of multistability in resting-state cortical rhythms. Journal of Neuroscience 31, 6353 (2011).

75. F. Freyer, J. A. Roberts, P. Ritter, and M. Breakspear. A canonical model of multistability and scale-invariance in biological systems. PLoS computational biology 8 (2012).

76. M. J. Aburn, C. Holmes, J. A. Roberts, T. W. Boonstra, and M. Breakspear. Critical fluctuations in cortical models near instability. Frontiers in physiology 3, 331 (2012).

77. D.-P. Yang and P. Robinson. Critical dynamics of hopf bifurcations in the corticothalamic system: Transitions from normal arousal states to epileptic seizures. Physical Review E 95, 042410 (2017).

78. M. J. Aburn. Critical fluctuations and coupling of stochastic neural mass models (2017).

79. K. J. Friston, A. Mechelli, R. Turner, and C. J. Price. Nonlinear responses in fmri: the balloon model, volterra kernels, and other hemodynamics. NeuroImage 12, 466 (2000).

80. E. M. Hillman, A. Devor, M. B. Bouchard, et al. Depth-resolved optical imaging and microscopy of vascular compartment dynamics during somatosensory stimulation. Neuroimage 35, 89 (2007).

81. D. Ress, J. K. Thompson, B. Rokers, R. Khan, and A. C. Huk. A model for transient oxygen delivery in cerebral cortex. Frontiers in neuroenergetics 1, 3 (2009).

82. P. Drysdale, J. Huber, P. Robinson, and K. Aquino. Spatiotemporal bold dynamics from a poroelastic hemodynamic model. Journal of theoretical biology 265, 524 (2010).

83. K. M. Aquino, P. A. Robinson, M. M. Schira, and M. Breakspear. Deconvolution of neural dynamics from fmri data using a spatiotemporal hemodynamic response function. Neuroimage 94, 203 (2014).

84. K. M. Aquino, M. M. Schira, P. A. Robinson, P. M. Drysdale, and M. Breakspear. Hemodynamic traveling waves in human visual cortex. PLoS Comput Biol 8, e1002435 (2012).

85. K. Aquino, P. Robinson, and P. Drysdale. Spatiotemporal hemodynamic response functions derived from physiology. Journal of theoretical biology 347, 118 (2014).

86. R. M. Birn, J. B. Diamond, M. A. Smith, and P. A. Bandettini. Separating respiratory-variation-related fluctuations from neuronal-activity-related fluctuations in fmri. Neuroimage 31, 1536 (2006).

87. J. D. Power, K. A. Barnes, A. Z. Snyder, B. L. Schlaggar, and S. E. Petersen. Steps toward optimizing motion artifact removal in functional connectivity mri; a reply to carp. Neuroimage 76 (2013).

88. K. Murphy and M. D. Fox. Towards a consensus regarding global signal regression for resting state functional connectivity mri. Neuroimage 154, 169 (2017).

89. Z. S. Saad, S. J. Gotts, K. Murphy, et al. Trouble at rest: how correlation patterns and group differences become distorted after global signal regression. Brain connectivity 2, 25 (2012).

90. K. Murphy, R. M. Birn, D. A. Handwerker, T. B. Jones, and P. A. Bandettini. The impact of global signal regression on resting state correlations: are anti-correlated networks introduced? Neuroimage 44, 893 (2009).

91. M. Allen, D. Poggiali, K. Whitaker, T. R. Marshall, and R. A. Kievit. Raincloud plots: a multi-platform tool for robust data visualization. Wellcome open research 4 (2019).

92. K. J. Friston, J. Kahan, B. Biswal, and A. Razi. A dcm for resting state fmri. Neuroimage 94, 396 (2014).

93. J. M. Shine. Neuromodulatory influences on integration and segregation in the brain. Trends in cognitive sciences 23, 572 (2019).

94. M. D. Fox, D. Zhang, A. Z. Snyder, and M. E. Raichle. The Global Signal and Observed Anticorrelated Resting State Brain Networks. J. Neurophysiol. 101, 3270 (2009).

95. A. Fornito, A. Zalesky, and E. T. Bullmore. Network scaling effects in graph analytic studies of human resting-state fmri data. Frontiers in systems neuroscience 4, 22 (2010).

96. A. Zalesky, A. Fornito, I. H. Harding, et al. Whole-brain anatomical networks: does the choice of nodes matter? Neuroimage 50, 970 (2010).

97. W. D. Penny, K. E. Stephan, A. Mechelli, and K. J. Friston. Comparing dynamic causal models. Neuroimage 22, 1157 (2004).

98. S. Oldham, A. Arnatkeviciute, R. E. Smith, et al. The efficacy of different preprocessing steps in reducing motion-related confounds in diffusion mri connectomics. Neuroimage In Press, 117252 (2020).

99. T. Obata, T. T. Liu, K. L. Miller, et al. Discrepancies between bold and flow dynamics in primary and supplementary motor areas: application of the balloon model to the interpretation of bold transients. NeuroImage 21, 144 (2004).

